# Neuroendocrine-like/EMT dedifferentiation mediates resistance to EGFR inhibitors via the NRG1/HER3 axis

**DOI:** 10.64898/2026.05.09.720556

**Authors:** Alessandra Morselli, Chiara Miroglio, William Kothalawala, Idan Lahat, Donatella Romaniello, Cinzia Girone, Francesca Ambrosi, Boobash Raj Selvadurai, Suvendu Giri, Michela Sgarzi, Martina Mazzeschi, Sabrina Valente, Andrea De Giglio, Gianandrea Pasquinelli, Arianna Palladini, Pier-Luigi Lollini, Michelangelo Fiorentino, Yosef Yarden, Yaara Oren, Balázs Győrffy, Andrea Ardizzoni, Mattia Lauriola

## Abstract

Patients with non-small cell lung cancer (NSCLC) carrying activating EGFR mutations typically respond favorably to third-generation EGFR tyrosine kinase inhibitors (TKIs) such as osimertinib. Nevertheless, resistance almost inevitably emerges, ultimately limiting the durability of these treatments. We investigated non-genomic mechanisms enabling drug-tolerant persister cells to survive EGFR inhibition, likely co-opting compensatory HER3 activation, whose underlying mechanisms remain unclear. Using a combination of immortalized and patient-derived cellular models, together with single-cell RNA sequencing, we demonstrate that activation of EGFR/HER3 axis constitutes an early adaptive response to TKI exposure enriched in pulmonary alveolar type I and type II cancer cells. This response is driven by neuregulin-1 (NRG1), produced by stromal cells and by cancer cells undergoing NE-like/EMT dedifferentiation. Importantly, *in vivo* studies demonstrated that combining EGFR inhibition and NRG1 neutralization by monoclonal antibody successfully eradicated tumors. Together, these findings point to a therapeutic strategy to overcome TKI resistance in NSCLC through targeting HER3 signalling, its interplay with EGFR, and tumor microenvironment–derived cues.

## Introduction

Oncogenic addiction is currently targeted primarily by tyrosine kinase inhibitors (TKIs) and monoclonal antibodies (mAbs), but the emergence of resistance limits the efficacy of both these agents. Non-small cell lung cancer (NSCLC) patients harboring activating EGFR mutations (deletion 746-750 in exon 19 and L858R mutation in exon 21) are treated with third-generation TKIs, such as osimertinib (*1*). Despite an initial positive response, patients develop resistance. Genomic alterations that underlie resistance to EGFR TKIs, such as the *EGFR* C797S mutation, or mutations in *MET*, *PIK3CA*, and *BRAF*, can now be routinely identified in clinical practice through genomic re-evaluation of patient samples, and are closely associated with the emergence of a resistant phenotype (*2*).

Before the development of genetically driven resistance, cancer cells often engage an early adaptive program, entering a drug-tolerant persister (DTP) state that is inherently difficult to capture. These cells exist in transient, reversible conditions that enable survival under therapeutic pressure through non-genetic tolerance mechanisms (*3*). One way for cancer cells to persist upon the treatment is through the activation of non-genomic signalling pathways. This involves triggering alternative signaling routes or bypass pathways. For example, one such pathway is the upregulation of HER3. In this case, cancer cells produce more RNA for the HER3 receptor, which allows to bypass the blocked EGFR and continue growing (*4–6*). Notably genomic alterations in HER3 are not known to constitute a mechanism of resistance to EGFR TKIs in EGFR-mutated NSCLC, while aberrant HER3 expression is associated with poor clinical outcomes in several tumour types(*7*). The relevance of this pathway has been the subject of an extensive debate, owing to the disappointing clinical outcomes obtained predominantly when monoclonal antibodies were employed as single agents. Only recently have the positive clinical results with the anti-HER3 antibody–drug conjugate patritumab deruxtecan provided clearer insight into the functional relevance of this pathway(*7*). This therapy was successfully applied in the HERTHENA trial with metastatic or locally advanced EGFR-mutated NSCLC, who had progressed on EGFR TKI therapy and platinum-based chemotherapy. The treatment showed clinically meaningful efficacy with increased PFS(*8–10*).

Nevertheless, the precise mechanism of action underlying HER3 dependency in patients who progress under TKI therapy remains unclear. In this study, the integration of immortalized and patient-derived cell models with single-cell sequencing technologies revealed a previously unrecognized symbiotic relationship between histologically distinct cancer clones. We found that, following TKI inhibition, HER3 becomes selectively enriched in DTP cells exhibiting pulmonary alveolar type I and II features. This dependency is further reinforced by paracrine signals, most notably NRG1, secreted by a subset of DTPs that undergo a neuroendocrine-like (NE-like) dedifferentiation program, as indicated by single-cell transcriptomic markers and the presence of electron-dense granules observed by electron microscopy.

NE-like dedifferentiated DTPs boost NRG1 production, which in turn impairs OSI response and enhances invasive phenotype, thereby prompting metastasis.

Notably, the combination of an antibody targeting NRG1, alongside a full neutralization of the EGFR pathway by a combo of anti-EGFR antibody and TKI cured mice in preclinical models.

## Results

### HER3 activation by NRG1 represents a bypass pathway in osimertinib-resistant patients

HER3 upregulation was previously reported as a bypass pathway promoting tolerance to EGFR inhibition. First, by treating with OSI the NSCLC cell line PC9, which harbours the founder EGFR driver *ex19* deletion, we confirmed the immediate downregulation of phospho-EGFR, associated with secondary upregulation of HER2 and HER3 in the population of drug tolerant persister cells (DTPs), which survived the high dose treatment *in vitro* (Fig. 1A). This led to a delayed activation of ERK and AKT, consistent with previously published data(*4–6*). In contrast, the downregulation of the MET receptor, as well as EGFR, persisted over time. We excluded the possibility that the upregulation of HER3 in DTPs was due to mere receptor stabilization, as mRNA levels were already increased after 24 hours of treatment (Fig. 1B). In contrast, EGFR expression showed a marked reduction, also at the mRNA level (Fig. 1B). Upon OSI treatment, the small fraction of DTPs began to exhibit high levels of NRG1, as confirmed by protein secretion into the medium, detected by ELISA as early as 24 hours after treatment (Fig. 1C). This may explain the delayed phosphorylation of HER3 previously detected. Since these results suggest that the bypass activation of HER3, associated with the acquisition of a DTP state, was driven by NRG1 production and secretion, we sought to investigate the role of NRG1 in the context of EGFR inhibition. To this end, we performed a 3D *in vitro* experiment and demonstrated that NRG1 administration was sufficient to impair the activity of OSI. Indeed, under NRG1 stimuli, OSI failed to induce cell death in otherwise sensitive cells (Fig. 1D). Interestingly, by re-analysing the data from Engelman and colleagues (*11*) which involved large-scale secretome profiling of NSCLC-stroma cultures, we found that NRG1 is highly expressed among NSCLC patient-derived fibroblasts treated with TKI (Fig. 1E). This suggests that its production in the microenvironment may influence the response to OSI.

**Fig. 1.**
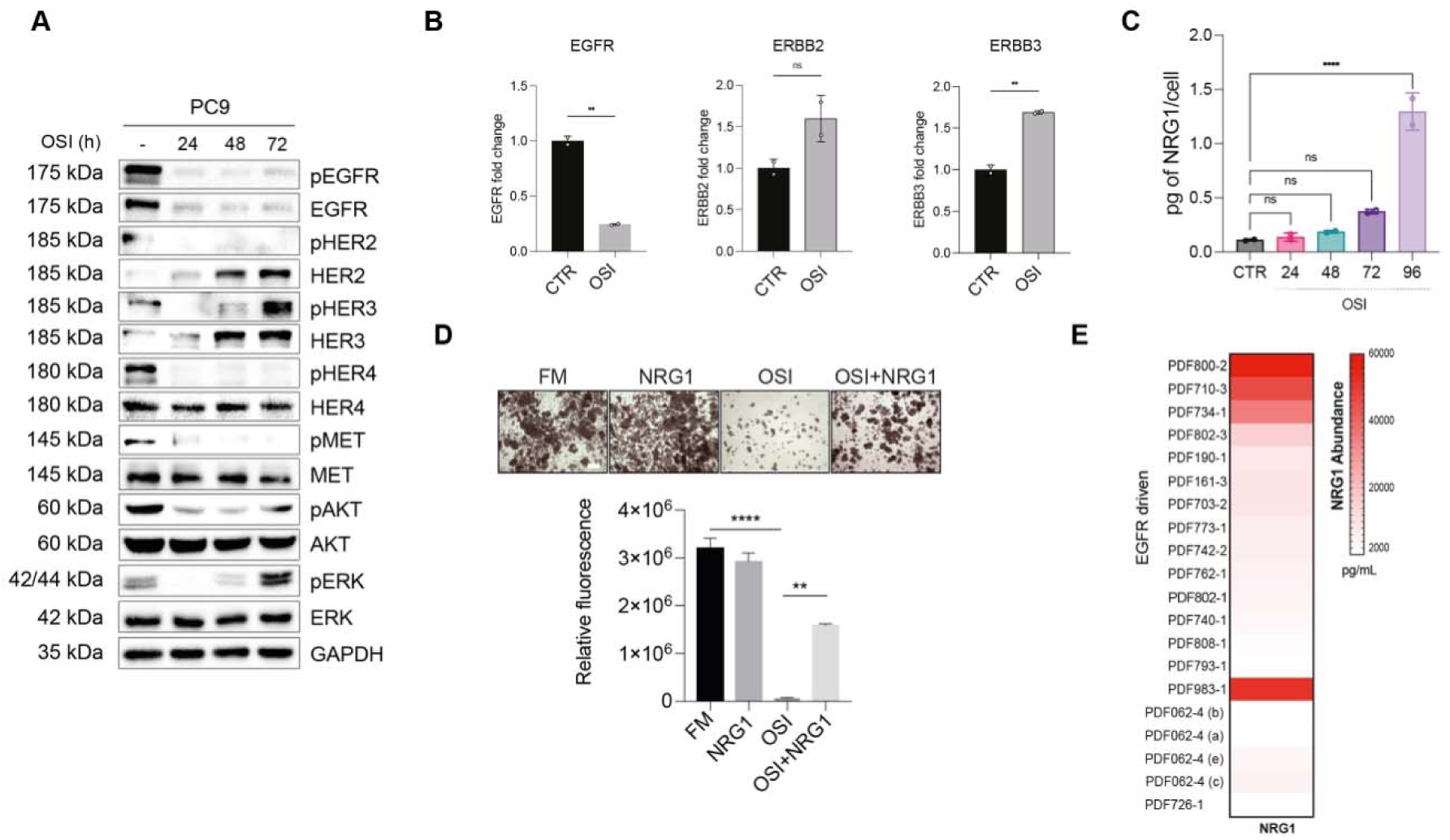
TKI treatment results in ERBB3 upregulation and NRG1 production: (**A**) Western Blot analysis on PC9 cells treated at the indicated hours with osimertinib (40 nM). Cell extracts were blotted and probed for the indicated proteins. GAPDH was used as a loading control protein. (**B**) PC9 cells were treated for 24 hours as in A. RNA was extracted and RT-qPCR was performed to analyze transcript levels of EGFR, ERBB2 and ERBB3. B2M was used as housekeeping gene for normalization. Graph reports mean ± SD and statistic was calculated by T-test. (**C**) PC9 cells were treated with OSI (40nM) and the secreted NRG1 in the conditioned media was measured using ELISA assay. After the treament cells were harvested, counted and used for data normalization. Graph bar reports mean ± SD and statistics was calculated by one-way ANOVA. (**D**) Spheroid-forming assays of PC9 cell line treated with OSI 40nM and NRG1 10ng/mL. Graph bar reports mean ± SD and statistics was calculated by one-way ANOVA. Scale bar: 200 μm. (**E**) Heat Map of NRG1 levels evaluated in the secretome of 20 patient derived fibroblasts. The NSCLC samples were all EGFR mutated and resistant to TKIs therapy. The bar shows the normalized protein expression (pg/ml per 100,000 cells).

### Establishment of ex vivo patient derived DTP cycling cells

The malignant pleural effusion from a patient receiving OSI treatment who continued to progress despite the absence of additional known mutations, was isolated, cultured *in vitro*, and designated as ADK11LM. Since these cells survived the OSI treatment in vivo, but regained sensitivity in vitro, we refer to them as early-DTPs (eDTPs), to distinguish from the sustained cycling-DTPs (cDTPs), which were generated upon further exposure to the drug *ex-vivo*. By simply propagating these cells *in vitro*, the immune cell population was eliminated, allowing the expansion of eDTPs cells capable of restoring their growth upon attachment to the culture plate (Fig. 2A).

**Fig. 2.**
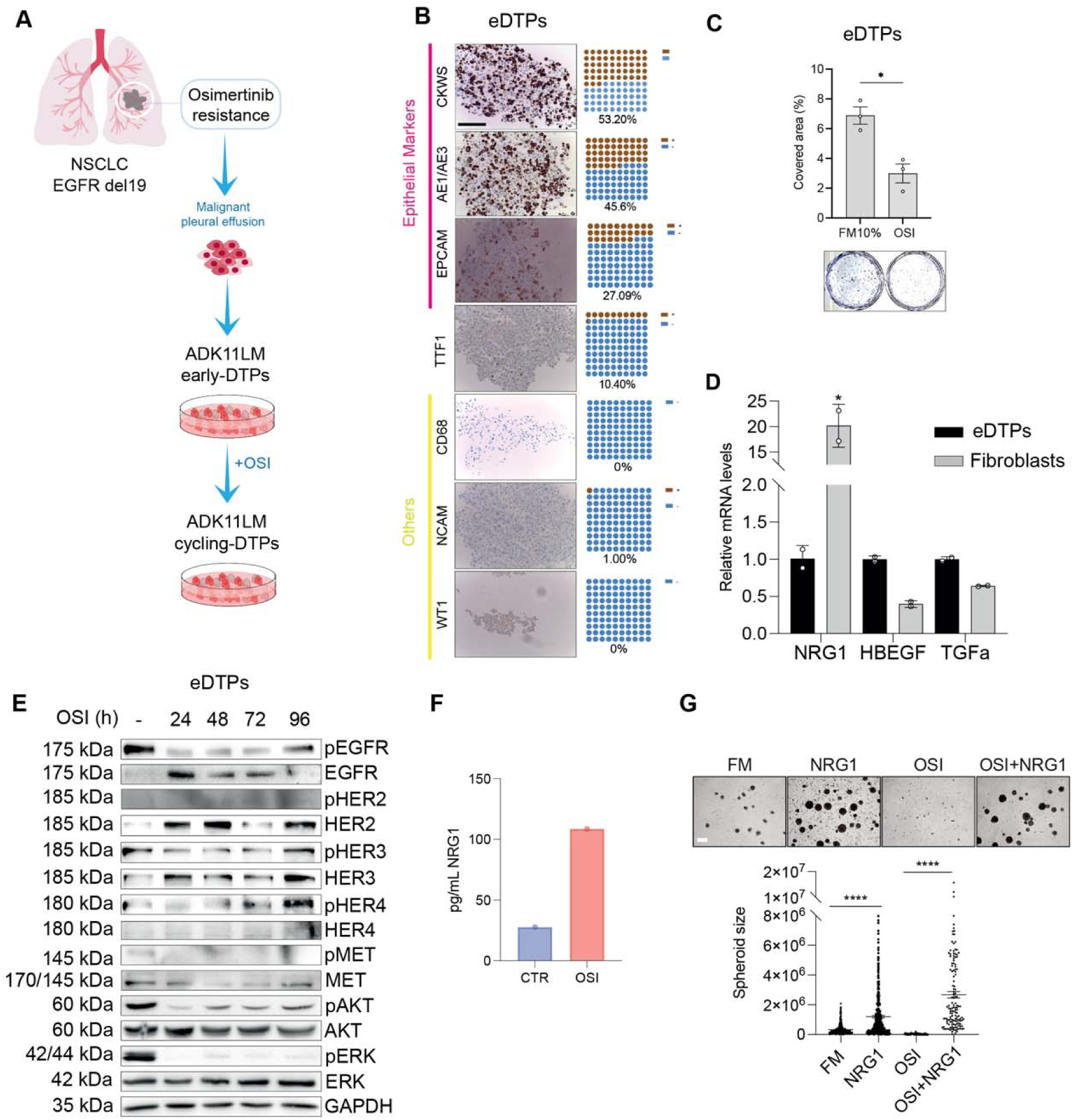
Characterization of patient derived DTPs: (**A**) Graphical model of ADK11LM early - DTPs tumor sample. Pleural effusion from a patient harboring an EGFR exon 19 deletion (ex19Del), who exhibited disease progression under OSI was cultured in vitro. Patient-derived tumor cells were then cultured with OSI in the media to obtain the corresponding cycling-DTPs. (**B**) IHC on early - DTPs tumor sample. The specimen was formalin-fixed and paraffin-embedded. We display respectively CKWS, AE1/AE3, EpCAM, TTF-1, CD8, NCAM and WT1 IHC (20X magnification). Scale bar: 100 µm. (**C**) Colony formation assay of early - DTPs tumor sample treated with OSI (40nM). Data are reported as % of wells covered area in relation to controls ± SD. Representative images of untreated and treated cells are shown. The statistic was calculated by unpaired T-test. (**D**) RNA was extracted from early - DTP tumor cells and fibroblasts derived from the same tumor sample separated through magnetic beads enrichment. RT-qPCR was performed to analyze transcript levels of NRG1, HBEGF and TGFa. Data are reported as fold changes related to control ± SD. Unpaired T-test was applied for statistical analysis. (**E**) Western Blot analysis on early - DTP cells treated at the indicated hours with osimertinib (40 nM). Cell extracts were blotted and probed for the indicated proteins. GAPDH was used as a loading control protein. (**F**) early - DTPs were treated with osimetinib (40nM) and the secreted NRG1 in the conditioned media was measured using ELISA assay. (**G**) Spheroid-forming assays of early - DTPs treated with OSI 40nM and NRG1 10ng/mL. Statistic was calculated by one-way ANOVA. Scale bar: 200 μm

Genomic profiles with the Oncomine Precision Assay (OPA) panel confirmed the presence of *ex19* deletion, and the lack of any additional mutations on known resistance pathways (table S1). Immunohistochemical (IHC) analysis of the eDTPs ADK11LM revealed that approximately 53% of the isolated cells were positive for pan-cytokeratin (panCK), indicating their epithelial origin (Fig. 2B). However, only 27% of these cells showed strong expression of epithelial cell adhesion molecule (EpCAM), suggesting a partial loss of epithelial characteristics and a dedifferentiation process already undergoing in this sample. Furthermore, the detection of thyroid transcription factor-1 (TTF-1) positivity in approximately 10.4% of the cells confirmed both the pulmonary adenocarcinoma nature of the tumor and its origin from alveolar type II cells (Fig. 2B). Through magnetic beads enrichment, we sorted patient-derived tumor cells. The lineage of the cultures was assessed via immunofluorescence using anti-vimentin antibody and through RT-qPCR. In the isolated non tumor cells, we detected fibroblast markers, namely up-regulated levels of collagen type 1 α2 (COL1A2), which is a subunit of type I collagen heterotrimers, and increased vimentin levels (Fig. S1A and Fig. S1B), this led us to confirm the presence of patient derived fibroblasts (PDFs). Notably, eDTPs tumor cells enrichment *in vitro,* appeared to restore the sensitivity to the OSI treatment (40nM). Indeed, when purified eDTPs were cultured *in vitro* and the stroma cells were removed, eDTPs retained partial sensitivity to OSI (Fig. 2C) in contrast with the progression observed in the corresponding patient.

NRG1 mRNA levels and two EGFR ligands, HB-EGF and TGFa were evaluated by RT-PCR showing that OSI-treated PDFs strongly overexpress NRG1, consistent with Engelman dataset(*11*) (Fig. 2D). Next, upon treatment with OSI, we biochemically assessed the eDTPs culture and observed a rapid downregulation of phosphorylated EGFR, accompanied by a compensatory increase in HER2 and HER3 expression (Fig. 2E). Notably, we detected increased phosphorylation of HER3 and HER4, but not of HER2. The signaling shift culminated in secondary upregulation of phosphorylated AKT, while MET activation remained negligible (Fig. 2E). Moreover, NRG1 secretion confirmed to be high upon OSI treatment (Fig. 2F). Finally, also in this second model, the 3D experiment showed that NRG1 administration was sufficient to strongly impair the activity of OSI in eDTPs, clearly proving that under NRG1 treatment OSI fails to induce cell death in otherwise sensitive cells (Fig. 2G).

### Cycling DTPs display a phenotypic shift promoting NRG1-mediated cell invasion and HER3 activation

Genomic analysis of eDTPs and cDTPs excluded the acquisition of known secondary mutations which may suggest the insurgence of resistance to the target therapy (table S1), and most importantly, OSI removal could restore sensitivity to the drugs (Fig. S2, A and B). These led us to confidentially classify these clones as cDTPs. By IHC analysis the ADK11LM cDTPs revealed that 100% of the cells were positive for panCK. However, only 8% of the cells expressed high levels of EpCAM with a 92% of cells undergoing EpCAM decrease, suggesting the emergence of a plastic, less differentiated cellular state (Fig. 3A). Interestingly, TTF-1 was maintained in about 10% of the cells (Fig. 3A). Characterization of the ERBB family in cDTPs cells revealed upregulation of EGFR, HER2, HER3 and HER4 at both protein and mRNA levels (Fig. 3, B and C), and HER3 ligand NRG1 was strongly upregulated (Fig. 3D). NRG1 is well known to induce motility and invasion by remodeling ECM proteins through the regulation of matrix metalloproteinases(*12*). Here, we found that eDTPs acquired a strong migratory and invasive phenotype upon exogenous NRG1 stimulation. Instead, cDTPs exhibited enhanced random motility and a markedly increased velocity even in the absence of exogenous NRG1, likely due to the autonomous NRG1 secretion (Fig. 4, A and B). Accordingly, cDTPs were able to invade the fluorescent gelatin in degradation assays. Extensive areas of matrix degradation, resulting from the formation of invadopodia in the ventral region beneath the nucleus, were detected as dark spots visible in the green channel, and became more pronounced upon NRG1 administration (Fig. 4C). These results may suggest that HER2/HER3 axis activation by NRG1 produces a progressive increase in terms of invasion in eDTPs. Consistently with an autocrine NRG1 production, cDTPs showed an enhanced migratory phenotype when compared to eDTPs, which is increased in both cell lines under NRG1 stimulation (Fig. 4C). Finally, by means of a monoclonal antibody neutralizing NRG1, we abolished gelatin degradation both with and without NRG1 administration (Fig. 4D), confirming the key role of NRG1 in cell invasion and metastasis of DTPs.

**Fig. 3.**
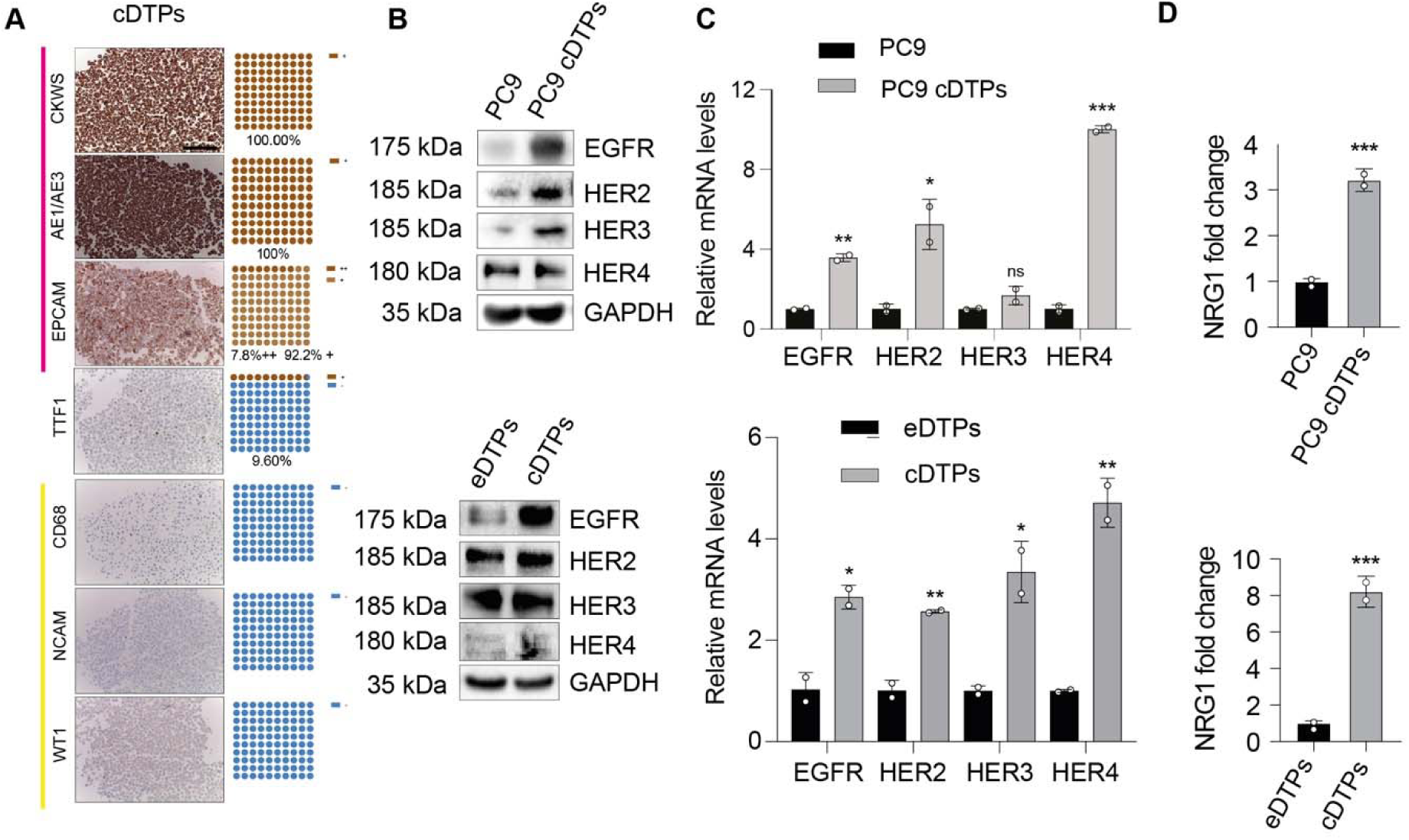
Establishment of patient derived cDTPs: (**A**) IHC on ADK11LM cycling-DTPs. All specimens were formalin-fixed and paraffin-embedded. We display respectively CKWS, AE1/AE3, EpCAM, TTF-1, CD68, NCAM and WT1 IHC (20X magnification). Scale bar: 100 µm. (**B**) Western Blot analysis on PC9 and the respective cycling DTPs (top panel) at the baseline state. The panel below shows the same analysis on ADK11LM early-DTPs and the respective cycling DTPs. GAPDH was used as loading control. (**C**) RNA was extracted from cell lines in (**B**) at the baseline state and RT-qPCR was performed to analyze transcript levels of EGFR, ERBB2, ERBB3, ERBB4. Graph reports mean ± SD and statistic was calculated by T-test. (**D**) Evaluation of NRG1 expression through qPCR in cell lines in (**B**). Graph reports mean ± SD and statistic was calculated by T-test.

**Fig. 4.**
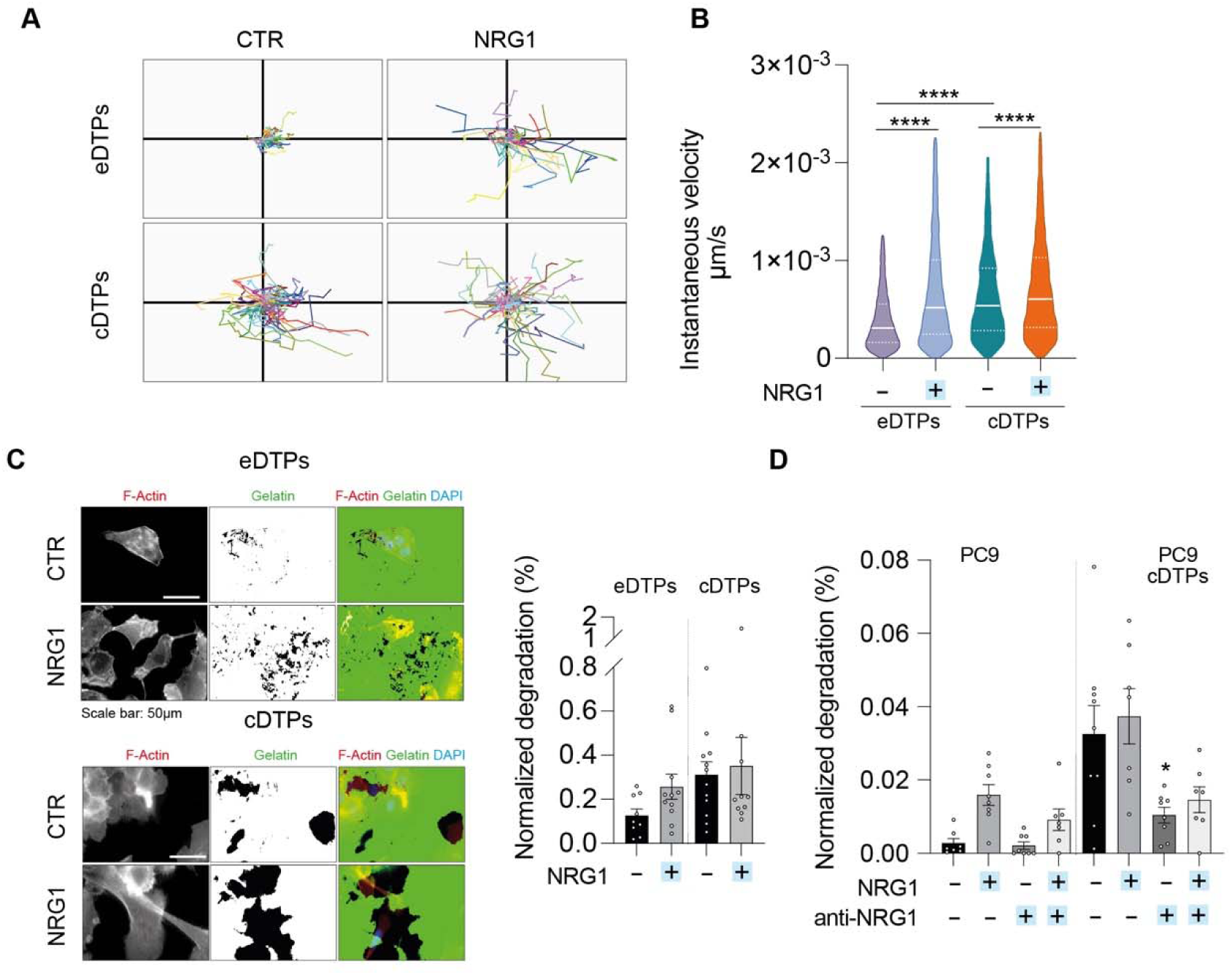
NRG1 prompts cell invasion and migration: (**A**) Cell tracking of ADK11LM early-and cycling-DTP cells stimulated with NRG1 10ng/mL and monitored for 48h with Phasefocus LiveCyte^TM^ platform. A total of 100 cell tracks are reported. (**B**) Charts showing the instantaneous velocity of cells treated and monitored over time as in **a**. Median and quartiles distribution are reported. Statistic was calculated by one-way ANOVA on cleaned data according to ROUT method (Q = 1%). 1500 cells were analyzed. (**C**) Gelatin-degradation assay performed on ADK11LM early- and cycling-DTPs treated with NRG1 10 ng/mL. Cells were stained with Phalloidin-TRITC (F-Actin) and DAPI (nuclei) prior to visualization. Invasion was quantified in terms of area of degraded gelatin (black spots visible in the green channel) normalized over the nuclei count in the same field. At least 7 fields were analyzed for each condition. Results are reported as fold increase related to control. Scale bar: 50 μm. (**D**) Gelatin-degradation assay performed on PC9 early- and cycling-DTP cells treated with NRG1 10 ng/mL and an anti-NRG1 mAb (YW538.24.71) 20 ug/mL. Cells were stained and then invasion was quantified as in **c**. Statistic was calculated by one-way ANOVA.

### Single-cell RNA sequencing analysis reveals the plasticity of lung cancer tumor cells

To better understand the mechanisms underlying the lack of response to OSI in the patient-derived model, we conducted a single-cell RNA sequencing (scRNA-seq) analysis. This approach unveiled transcriptional variability at high resolution in both the primary eDTPs and the cDTPs. Briefly, cells isolated from each population were captured in droplet emulsions using a Chromium Single-Cell instrument (10x Genomics) and libraries were prepared and sequenced on a NovaSeq 6000 (Illumina). Through bioinformatic analyses of RNAseq data, we identified seven main clusters in eDTPs, confirming the intrinsic heterogeneity of this patient-derived sample, as already suggested by IHC marker characterization (Fig. 5A). First, we confirmed the presence of a stromal population, accounting for approximately 21.5% of the sample, identified as fibroblasts based on the expression of genes such as *COL3A1*, *COL5A2*, and *VIM*, and the positivity for VIM evaluated by immunofluorescence and IHC (Fig. S1, A, B and C). In addition, we identified several clusters of well-differentiated epithelial tumor cells falling within pulmonary alveolar type 1 (AT1) cells (13%), alveolar type 2 (AT2) cells (10%) which are highly differentiated cells lining the lung alveoli, and Clara cells (7%) located in the terminal duct. In contrast, we observed a distinct cluster of *KRT*-positive cells that had lost their lineage-specific differentiation markers. Although these cells could no longer be classified as AT1, AT2, or Clara cells, they retained features consistent with an epithelial phenotype, such as high expression of *KRT19*, *KRT18*, and *EPCAM*, as well as the presence of the *EGFR* exon 19 deletion mutation. We therefore classified this cluster simply as cancer cells (15%). Consistent with the notion that epithelial cells within tumor specimens represent a heterogeneous continuum of mutant subpopulations, we also identified a cluster of cells undergoing complete dedifferentiation. These cells were marked by downregulation of *EPCAM* and upregulation of epithelial-to-mesenchymal transition (EMT) markers, including the mesenchymal regulators *TWIST1/2* and *ZEB1/2*, as well as genes associated with mesenchymal lineages such as *FBN1/2*, *CD44*, *COL5A2*, *COL6A2*, *COL4A2*, *COL16A1*, *COL5A1*, *COL7A1*, *COL12A1*, *COL6A3*, *CDH6*, *CDH11*, *FAP*, *PLOD2*, *POSTN*, and *VCAN*. Notably, markers of neuronal and neuroendocrine lineages, including *NCAM1*, *NCAM2*, *CDH2*, and *CDH4*, were also expressed. The downregulation of canonical epithelial markers in this cluster likely impaired accurate annotation by the automated bioinformatic tool, which failed to assign it to a known cancer category. Thus, we manually reviewed the associated gene expression profiles and integrated additional cell-type markers. Based on these features, we were prone to classify this cluster as NE-like/EMT dedifferentiated cells (25%) (Fig. 5A). Finally, a cluster of alveolar macrophages or dust cells (8%) was also detected (Fig. 5A). The overall single-cell transcriptomic data supported a model in which TKI response promotes diversification of cell state; thus, we performed the same analysis on cDTPs (Fig. 5B). Here, we identified only five cell clusters, as certain populations, such as stromal fibroblasts as well as alveolar macrophages were lost during *in vitro* passaging, likely due to terminal senescence. In contrast, the populations of AT-2 cells, AT-1 cells, and Clara cells steadily decreased, accounting for 10.6%, 10.2%, and 3.4% of the total population, respectively. Interestingly, the cancer cells cluster was now absent, and the *KRT7, KRT8, KRT10, KRT13, KRT16, KRT18, KRT19* positive cells were classified as basal cells (airway progenitor cells), suggesting that cancer cells were likely undergoing a shift in plasticity toward a less differentiated phenotype in cDTPs (9%). Finally, we unveiled a large, poorly defined cluster (approximately 67%) that exhibited features of NE-like/EMT dedifferentiation, including *TWIST1*, *SNAI3*, *MMP14*, *FBN1*, *CD44*, *COL3A1*, and *COL6A3* and the increased expression of neuronal and neuroendocrine lineage markers, pointing to a pronounced shift toward NE-like differentiation. Indeed, in addition to *NCAM2* and *CDH4*, we detected expression of *NEUROD1*, *SYPL2*, *PTH*, *NRCAM*, *DSCAM*, NRXN3, SNCA, and *OXTR*, highlighting the neuroendocrine drift within the tumor compartment in cDTPs. Going into the details, in eDTPs ERBB family expression was detected across all cell clusters, with particularly high levels observed in pulmonary AT-1 and AT-2 cells, Clara cells, cancer cells and very low level in the NE-like/EMT dedifferentiated cluster (Fig. 5C). *ERBB2* levels were especially elevated in AT-1 and AT-2 cells, while *ERBB3* and *ERBB4* showed very low expression across the population. In contrast *MET* and *AXL* were specifically upregulated in the NE-like/EMT population in the eDTPs (Fig. 5C). In contrast, in cDTPs, we found that *ERBB2* and *ERBB3* levels strongly increased in AT-1 and AT-2 cells, Clara cells and basal progenitor cells (Fig. 5D). Finally, *NRG1* appeared differentially distributed among the identified cell populations (Fig. 5, E and F). In the eDTPs, the major producers of NRG1 were fibroblasts, alveolar macrophages and dedifferentiating epithelial tumor cells including NE-like/EMT dedifferentiated cells (Fig. 5E). Interestingly, in the cDTPs, *NRG1* production shifted almost exclusively to the NE-like/EMT dedifferentiated cells (Fig. 5F). These findings suggest that, in eDTPs, the proliferation and survival of cancer cells may initially be supported by NRG1 secreted primarily by the stroma. The survival of cDTPs under OSI treatment may instead rely on NRG1 production by tumor cells themselves, particularly those undergoing NE-like/EMT transdifferentiated drift, thereby promoting continued proliferation and tolerance to the drug.

**Fig. 5.**
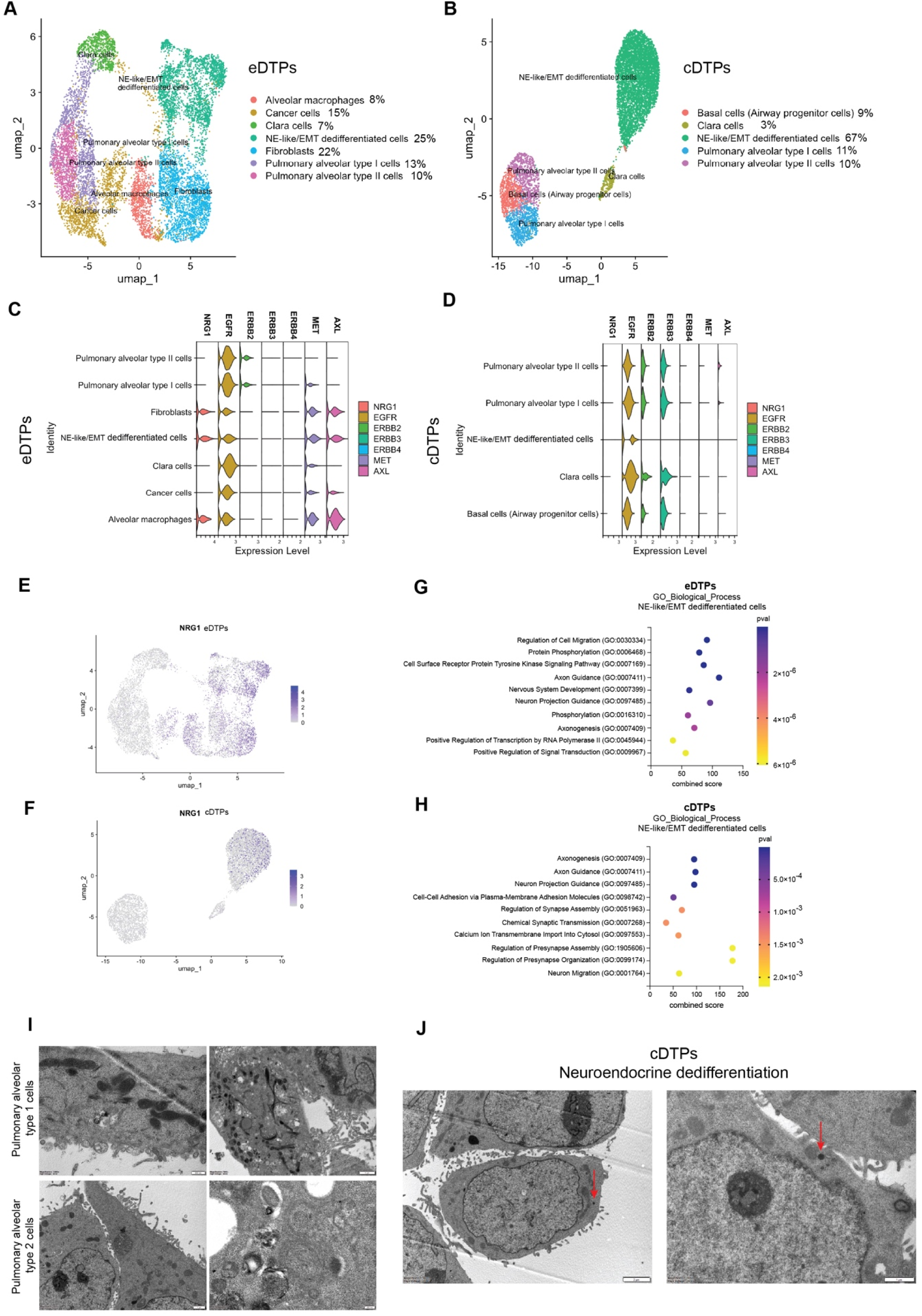
ScRNAseq on patient derived eDTPs and cDTPs: (**A** and **B**) UMAP plot of all cells clustered and color coded by cell type in ADK11LM early- (**A**) and cycling-DTPs (**B**) samples analyzed through scRNAseq. Clusters were assigned to the indicated cell types by differentially expressed genes (DEGs) and analyzed with ScType computational platform. (**C** and **D**) Violin plot shows distribution of RTKs expression across different clusters in (**C**) and cycling-DTPs (**D**). **e** UMAP plot of NRG1 mRNA expression across all cell types in ADK11LM early-DTP sample. (**F**) Same analysis reported in **e** but in ADK11LM cycling-DTPs. (**G** and **H**) Dot plot of pathway enrichment analysis performed on scRNAseq data showing the top 10 enriched GO BP with differential expression between the early and cycling - DTPs and the other cell types. The analysis was performed using the online tool Enrichr. Dot plot of pathway enrichment analysis performed on scRNAseq. Data showing the top 10 enriched Hallmark with differential expression between the early and cycling-DTPs and the other cell types. The analysis was performed using the online tool Enrichr. (**I**) Transmission electron microscopy (TEM) of early- and cycling-DTP cellular samples. The images shown AT1 and AT2 differentiative characteristics. (**J**)Transmission electron microscopy (TEM) of cycling-DTP cellular sample whose cytoplasm contains small electron-dense-cored granules (arrows). Each scale bars are reported on the images.

Next, we sought to further investigate the molecular pathways of the NE-like /EMT dedifferentiated cell clusters, which were not classified by standard single-cell annotation methods. To this end, we performed analysis on the top 1,000 differentially expressed genes (DEGs) to uncover gene signatures most enriched in this population. The top 10 statistically over-represented GO BP (biological process) terms and hallmark are shown in Fig. 5, G and H and Fig. S3, A and B. Interestingly, ENRICHR analysis of the eDTPs revealed significant enrichment in biological processes involved in the regulation of cell migration, axon guidance, nervous system development, neuroprojection guidance as well as axonogenesis, suggesting neuronal differentiation programs (Fig. 5G). Similarly, the NE-like/EMT dedifferentiated cluster of cDTPs showed even stronger enrichment, with a smaller p-value, for biological processes related to axonogenesis, axon guidance, and neuron projection guidance (Fig. 5H). This finding is consistent with the increase in the proportion of cells belonging to this cluster, which rose from 25% in the eDTPs to over 60% in the cDTPs. These results increased our confidence in classifying these cells as undergoing differentiation toward an NE-like/EMT trajectory. Due to the lack of well-established markers for NE-like cells, we performed ultrastructural microscopy analysis on these samples, which revealed the presence of distinct features. Specifically, we observed partially flattened cells containing bundles of cytokeratin filaments and vacuoles, typical of the AT1 morphology (Fig. 5I). In addition, cuboidal cells with well-developed lamellar bodies were identified, matching the histological features of AT2 cells (Fig. 5I). Finally, we observed a population of cells containing cytosolic electron-dense granules (Fig. 5J). These granules may corroborate the single-cell-based gene expression annotation and further support the conclusion of a bona fide NE-like dedifferentiation drift occurring at an early stage of DTPs, detectable by ultrastructural microscopy and not yet revealed by canonical markers such as SYP or NCAM by IHC. Next, to further investigate the plasticity of the NE-like/EMT module we unveiled, we examined its abundance in DTPs over time. Strikingly, NE-like/EMT module score showed a progressive and steady increase during the OSI course of treatment in PC9 cells, becoming evident as early as day 3. Notably, its enrichment was more pronounced in the non-cycling DTP after 14 days of treatment, suggesting that acquisition of NE-like/EMT features may preferentially occur in a quiescent drug-tolerant state (Fig. 6A). We next evaluated NRG1 expression in patients using scRNA-seq dataset from Bivona et al(*13*). NRG1 expression was undetectable in treatment-naive samples, increased in persister cells, and reduced at relapse while remaining above naive levels (Fig. 6B), consistently, in patient TH226, which had matched biopsies from different time points, NRG1 expression was absent in the treatment-naive sample, increased in persisters during ongoing OSI treatment, and decreased at relapse (Fig. S3C). We then examined whether *NRG1* expression tracked with NE-like/EMT transcriptional state changes across the tumor and microenvironment. While the NE-like/EMT-score did not show a global increase in patient-derived data, a small subset of cells displays NE-like/EMT-score higher than the naïve baseline (Fig. S3D). Notably, positive *NRG1*– NE-like/EMT correlations were observed specifically in the persister stage and were restricted to tumor cells and non-tumor fibroblasts, suggesting that these rare NE-like/EMT-high cells may be responsible for driving the correlation observed in Fig. 6C. In contrast, relapse samples showed no correlation (Fig. 6C). Consistent with this pattern, persister tumor cells showed a clear positive relationship between NRG1 expression and NE-like/EMT-like module score (Fig. S3E). Finally, we implemented further analyses of real-life samples collected from patients with NSCLC (Fig. S4A) undergoing TKI treatment over time (on-treatment) and after disease progression. Notably, one patient progressing with OSI exhibited a pleomorphic transition associated with MYC amplification. Histological analysis revealed a bipartite tumor, comprising a component resembling the pre-treatment biopsy and an additional NE-like/EMT dedifferentiated carcinoma component consistent with pleomorphic carcinoma (Fig. S4B). Molecular analysis confirmed the upregulation of *ERBB3* or *NRG1* transcript levels (Fig. S4, C and D). Moreover, *ASCL2*, *SYPL1* and *NRG1* were found to be upregulated in the pleomorphic component supporting the acquisition of a NE-like/EMT transcriptional program (Fig. S4D). Altogether, these results underscore a functional association between NRG1 and NE-like/EMT features in drug-tolerant states, supporting the clinical significance of these findings.

**Fig. 6.**
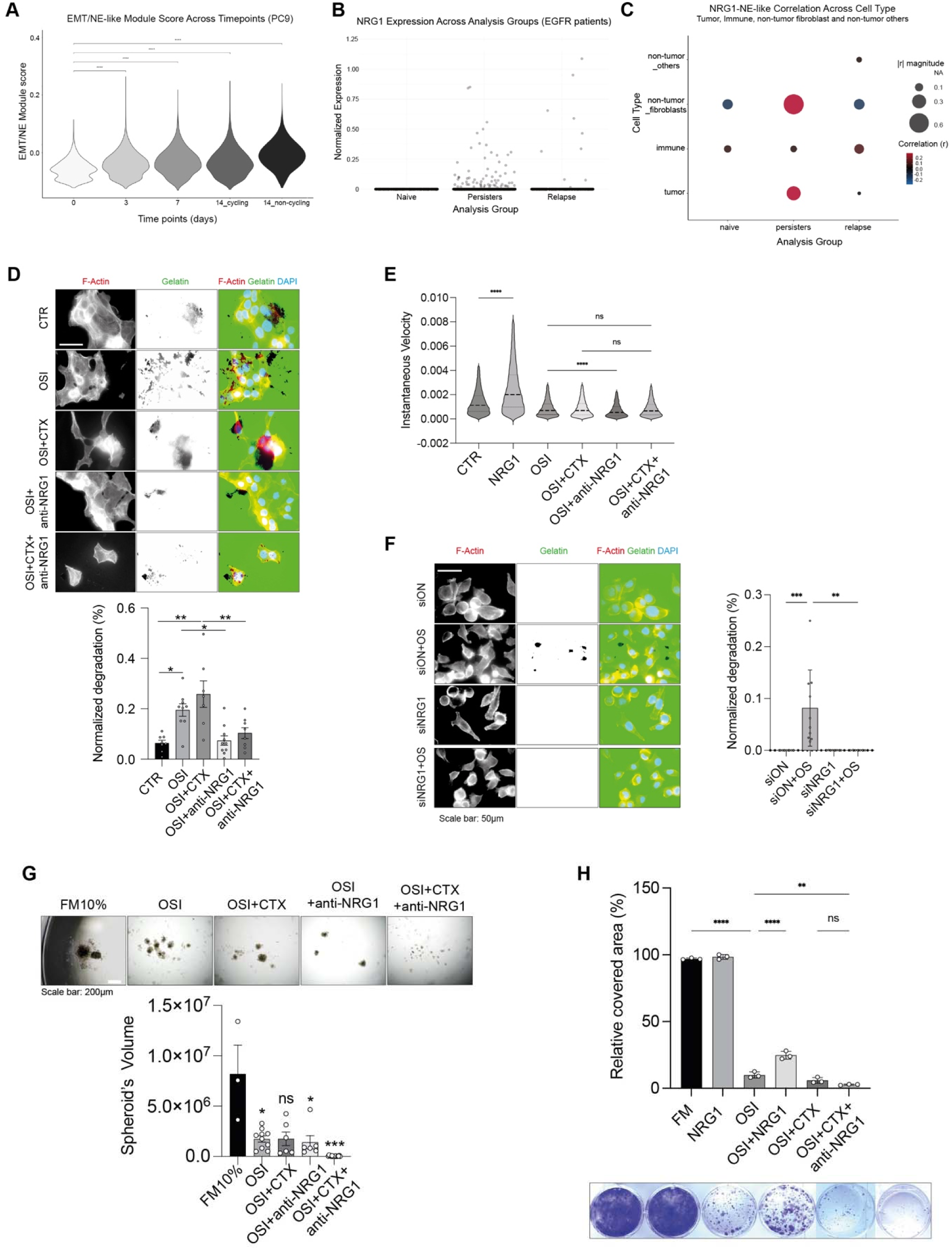
Upregulation of NRG1 in EMT-NE–like persister cells promotes invasion and tolerance to osimertinib: (**A**) EMT/NE-like module score in PC9 cells across osimertinib time course (days 0, 3, 7, and day 14 cycling vs non-cycling DTPs). Statistics were calculated by pairwise Wilcoxon rank-sum test versus day 0 and significance is indicated by stars. (**B**) NRG1 expression across EGFR-mutated NSCLC patients comparing naive, persisters, and relapse samples (Bivona dataset). Each dot represents a single cell. (**C**) Pearson correlation between per-cell NRG1 expression and EMT/NE-like module score across clinical groups and cell types. (**D**) Gelatin-degradation assay performed on early-DTPs sample. The following treatments were applied: osimertinib (40nM), osimertinib + cetuximab (20 ug/mL), osimertinib + anti-NRG1 mAb (20 ug/mL) and the triple combination. When combined the doses of the two mAb were halved. After 24h, cells were stained with Phalloidin-TRITC (F-Actin) and DAPI (nuclei) prior to visualization. Invasion was quantified in terms of area of degraded gelatin (black spots visible in the green channel) normalized over the nuclei count in the same field. At least 7 fields were analyzed for each condition. Scale bar: 50 μm. Statistic was calculated by one-way ANOVA. (**E**) Quantification of ADK11LM early-DTPs instantaneous velocity over a 72h time-lapse obtained with the Phasefocus LiveCyte™ system, capturing images every 2h. Median and quartiles distribution are plotted. Outliers were identified and cleaned from results by means of ROUT method (Q = 1%). Statistic was calculated by one-way ANOVA test. (**F**) Gelatin-degradation assay performed on PC9 cells with a non-targeting control siRNA (siON) or siRNA targeting NRG1 (siNRG1). After siRNA transfection, cells were seeded and after 6 hours treated with osimertinib (40nM). After 24h, cells were stained with Phalloidin-TRITC (F-Actin) and DAPI (nuclei) prior to visualization. Invasion was quantified in terms of area of degraded gelatin (black spots visible in the green channel) normalized over the nuclei count in the same field. Scale bar: 50 μm. One-way ANOVA test was applied. (**G**) Spheroid-forming assay on early-DTPs treated with: osimertinib (40nM), osimertinib + cetuximab (20 ug/mL), osimertinib + anti-NRG1 mAb (20 ug/mL) and the triple combination. When combined the doses of the two mAb were halved. Spheroid-forming ability was measured by spheroids size calculated through the formula (minor axis x major axis2) x 2. The statistic was calculated by one-way ANOVA. Scale bar: 200µm. (**H**) Colony formation assay of PC9 cells treated with NRG1 (10 ng/mL), osimertinib (40nM), osimertinib + cetuximab (20 ug/mL), osimertinib + anti-NRG1 mAb (20 ug/mL) and the triple combination. When combined the doses of the two mAb were halved. Cells were seeded in triplicate in 12-well plates and treated after 24 hours. Following a 10-day incubation period, cells were fixed using 4% PFA and stained with crystal violet for 30 minutes. Colony formation ability was quantified by counting the colonies using ImageJ software. Data are reported as % of wells covered area in relation to controls ± SD. The statistic was calculated by one-way ANOVA test.

### NRG1 neutralization nullifies invasion and restores OSI effectiveness

As revealed by both single-cell RNA sequencing data and complementary *in vitro* experiments, NRG1 secreted by the NE-like/EMT dedifferentiated cells emerged as a pivotal mediator driving tolerance to EGFR inhibition. To further explore this mechanism and evaluate the therapeutic potential of targeting NRG1, we employed a specific neutralizing antibody against NRG1. First, we proved the ability of a mAb anti-NRG1 to reduce cell invasion in eDTPs, when combined with the combo OSI + CTX (Fig. 6D). Interestingly, while OSI alone or in combination with CTX enhanced the invasive capability, likely due to the activation of compensatory HER2 and HER3 pathways, the addition of antiNRG1 neutralizing antibody partially disrupted this effect, probably by impairing EGFR/HER2 or EGFR/HER3 dimers that may support invasion upon NRG1 secretion. Similar results were confirmed on cell velocity which represents a further proxy of the migration (Fig. 6E and Fig. S5, A, B, C and D). Next, to demonstrate mechanistically that DTPs invasion occurred through NRG1 secretion, we performed NRG1 siRNA silencing. First, we confirmed efficient NRG1 knockdown, with transcript levels reduced by approximately 80% (Fig. S5E). Strikingly, NRG1 silencing was sufficient to block eDTPs (Fig. 6F) and invading cDTPs (Fig. S5F).

The role of NRG1 neutralization in the tolerance to OSI was also tested. While DTPs managed to survive in 3D, the addition of an anti-NRG1 antibody combined with CTX completely abolished it, both in 3D culture (Fig. 6G) and in standard adherent conditions (Fig. 6H). In summary, these findings support a model in which NRG1 secretion enables the activation of EGFR/HER3 and EGFR/HER2 heterodimers, which sustain tolerance to OSI and the invasive potential. Remarkably, NRG1 neutralization interferes with HER2/HER3 signaling complexes, as a result, effectively suppresses DTPs survival and invasion.

### The anti-NRG1 mAb in combination with EGFR full neutralization eradicates tumors and prevents RTKs activation

Finally, to further confirm the preclinical relevance of these *in vitro* data, we tested the anti-NRG1 as a strategy to reduce the resistance to last-generation EGFR inhibitors in an *in vivo* setting. We employed 6-weeks old female athymic nude Foxn1nu mice, that were inoculated with 3×10^6^ PC9 cells in each animal’s right flank. Mice harboring palpable tumors were divided into five different treatment groups: vehicle control, OSI (5mg/kg), OSI+CTX (5mg/Kg + 0,2mg/mouse/injection), OSI+mAb anti-NRG1 (5mg/Kg+0,2mg/mouse/injection), OSI+CTX+mAb anti-NRG1(5mg/Kg + 0,1mg/mouse/injection for each antibody). Treatments were administered twice weekly intraperitoneally (CTX and mAb anti-NRG1) and by daily oral gavage (OSI) continued until day 24. Animals were then monitored for an additional month. Tumor volume was evaluated twice a week, along with measurement of the body weight (once a week). Mice were euthanized when tumor size reached 1,500 mm^3^. OSI monotherapy caused rapid, yet incomplete tumor shrinkage, followed by recurrence during the drug holiday (Fig. 7A). Similarly, neither of the two drug combinations, OSI + anti-NRG1 and OSI + CTX, fully prevented relapse, although the latter was clearly superior. In contrast, the triplet combination was curative in 5 mice out of 10: within 2–3 weeks all tumors disappeared, and none relapsed after we stopped the treatments. In addition, this combination proved the best results in terms of tumor growth and animal survival (Fig. 7, B and C). Notably, although body weight of all mice was measured once a week, we observed no significant deviation from the control group, indicating absence of toxic effects. In parallel, to resolve the molecular basis, we re-performed the animal trial with three mice per group sacrificed shortly after treatment onset (on day 7). Thereafter, tumors were processed for immunoblotting (Fig. 7D). Despite inter-animal variation, the results linked the therapeutic effects to the ability of anti-NRG1 to downregulate both the total levels of HER3, HER2 and MET as well as to prevent their activation. Consequently, the active forms of both AKT and ERK were severely reduced. Finally, we evaluated HER3 levels in the same tumors collected from mice by IHC, confirming the upregulation of HER3 upon OSI treatment (Fig. 7E). In conclusion, when combined with CTX and OSI, the anti-NRG1 mAb used, prevented tumor relapse long after all treatments were stopped, likely due to the simultaneous inactivation of several bypass pathways.

**Fig. 7.**
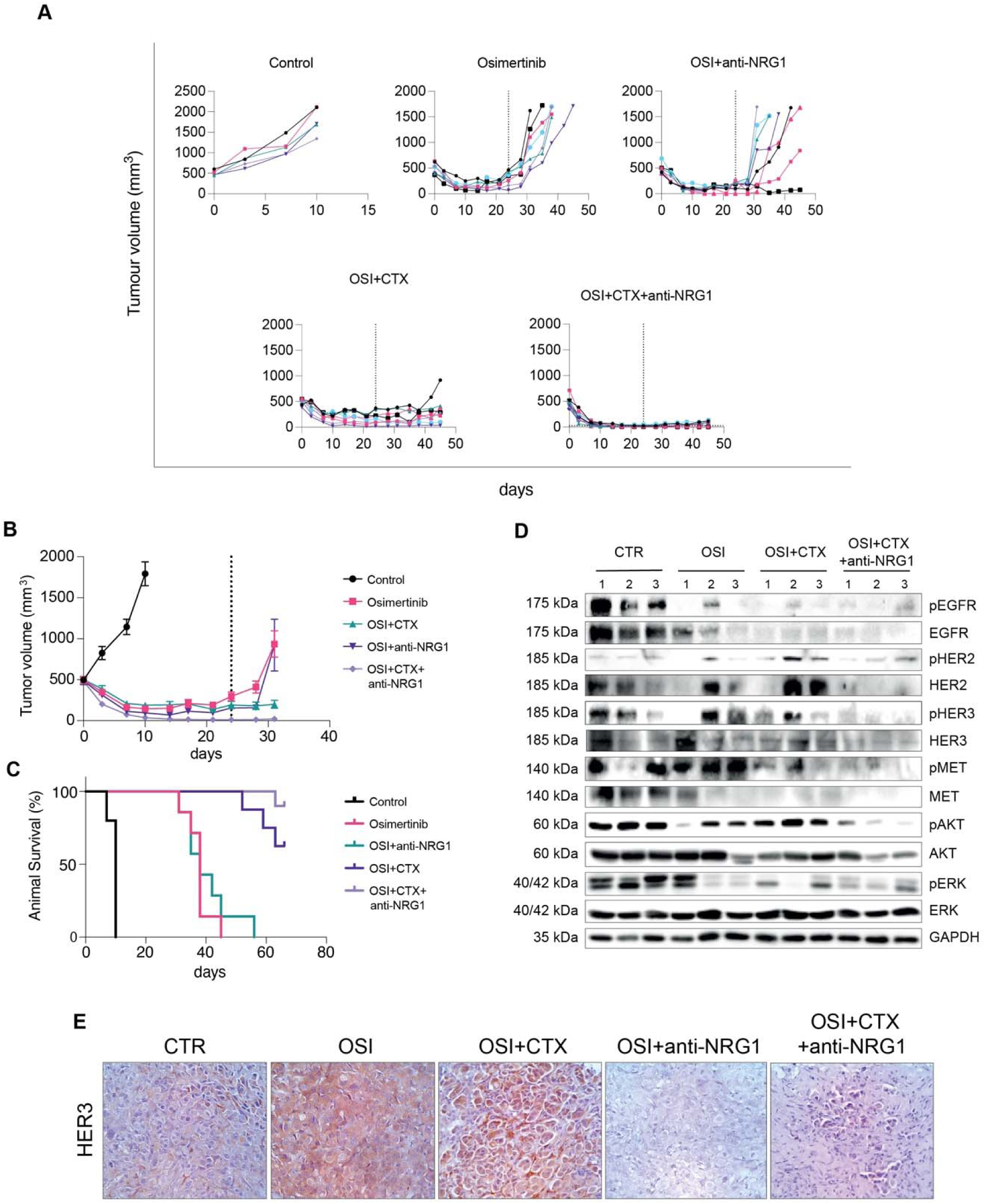
In vivo validation of NRG1 neutralization effectiveness in reducing resistance to EGFR-TKIs: (**A**) CD1 nu/nu mice were injected with PC9 cells (E746–A750 deletion; 3 × 10^6^ cells per mouse). Mice harboring palpable tumors were divided into five different treatment groups: (i) Vehicle control, (ii) daily oral gavage of Osimertinib (administered at 5 mg/kg/mouse), (iii) Osimertinib + cetuximab (5mg/Kg + 0,2mg/mouse/injection), (iv) Osimertinib + mAb anti-NRG1 (5mg/Kg + 0,2mg/mouse/injection), (v) Osimertinib + cetuximab + anti-NRG1 (5mg/Kg + 0,1mg/mouse/injection for each antibody). The antibodies were intraperitoneally injected twice a week. Treatments continued until day 24 and tumor volume was monitored until day 66. Mice were euthanized when tumor size reached 1500 mm3. (**B**) Tumorigenic growth over time of PC9 derived tumors. (**C**) Survival assessment of mice in A. (**D**) Cleared whole extracts of individual tumors were blotted and probed for the indicated proteins. GAPDH was used as loading control protein. (**E**) Immunohistochemical evaluation of HER3 expression in PC9 xenograft tumors extracted from mice.

## Discussion

Harnessing ERBB bypass pathways to overcome TKI inhibition of mutant EGFR is a well-established route of acquired resistance in NSCLC. Our findings help to shed light on a possible mechanism by which patients fail to respond to OSI. Indeed, we first demonstrated that TKI treatment in DTPs significantly enhances the secretion of the NRG1 ligand, which occurs in cells undergoing NE-like/EMT dedifferentiation. At the same time, TKI treatment induces the *de novo* synthesis of ERBB3, the principal binding partner of NRG1, within the well-differentiated compartment of the tumor bulk. Due to its biochemical nature, lacking an active kinase domain, ERBB3 is incapable of independently transducing downstream signals. As a result, activated ERBB3 is compelled to heterodimerize with other ERBB family members to initiate signal propagation. This is particularly concerning because our data indicate that DTPs that persist following treatment, despite inhibited proliferation, exhibit a marked propensity for gelatin degradation, a phenotype associated with increased invasive potential *in vivo*. Notably, we found that this invasive behaviour is dependent on HER3 heterodimers formation, indeed, NRG1 presence enhances this effect. Importantly, neutralization of NRG1, achieved either through siRNA-mediated silencing or the use of a neutralizing monoclonal antibody, completely abrogated tolerance as detected by lack of invasive phenotype and tumor growth *in vivo*.

Mechanistically, single-cell analysis of samples from patients non-responsive to OSI revealed that NRG1 secretion is predominantly sustained by stromal cells, including both fibroblasts and alveolar macrophages (*11*). Prolonged drug exposure induced plastic histological transformation into a dedifferentiated state of the cDTPs. First, in this work we observed a cellular heterogeneity within the DTPs, including distinct lung epithelial lineages such as AT-1 and AT-2 cells, as well as ductal (Clara) cells. Notably, the eDTPs presented an enrichment of the alveolar cell signature, consistent with the results previously reported in patients(*13*), and a tendency toward a decrease in this program in the sustained cDTPs. Intriguingly, a distinct cluster that expressed markers of NE-like/EMT differentiation emerged, likely as an adaptive response to drug-induced stress. The abundance of this NE-like/EMT population increased significantly following extended OSI treatment. This cluster displayed low expression of EGFR family members, which may account for tolerance to the TKI(*14*). Notably, the transition from eDTPs to cDTPs, was associated with global reduction of EGFR within the NE-like/EMT population, suggesting an adaptive regulation at single cell levels, rather than selection of a pre-existing population. This observation is fully consistent with the histological transformation reported in murine models of lung adenocarcinoma, in which chemical or genetic inhibition of EGFR induces a marked expansion of Ascl1-expressing cells, the definitive lineage marker of the neuroendocrine state.(*15*) Notably, the eDTPs population exhibited elevated AXL expression, a feature previously implicated in tolerance mechanisms(*16*)and in promoting a hyper-mutant phenotype, thereby contributing to the emergence of therapy resistance(*17*).

Finally, the more differentiated cancer cells maintained higher levels of EGFR, HER2, and HER3. Importantly, the NE-like/EMT cells mainly contributed to tumor progression by secreting NRG1, thereby activating the HER3/EGFR bypass pathway and supporting continued growth despite EGFR-targeted therapy. Consistently, NRG1/HER3 signalling in circulating tumour cells has been implicated in supporting CTC survival and facilitating metastatic dissemination in breast cancer(*18*). In line, in patients with advanced NRG1 fusion–positive cancers, such NSCLC and pancreatic cancer, the bispecific ERBB3/ERBB2 antibody led to impressive clinical responses, accompanied by low-grade adverse events(*19*). These findings support a role for NRG1 as a major oncogenic driver and suggest that its neutralization, either through monoclonal antibodies or ERBB3-directed bispecific antibodies as well as by HER2 TKI(*20*) may offer a promising treatment. NRG1 and ERBB2 axis activation has been implicated in resistance also to monoclonal antibody, targeting EGFR(*21, 22*). More broadly, paracrine NRG1 activation, originating from tumor-associated fibroblasts or adipocyte precursors, has been reported to promote antiandrogen resistance(*23*) or resistance to drugs targeting receptor tyrosine kinases such as FGFR or ALK(*24, 25*), through alternative HER3 activation pathways. Considering that patient-derived cells display nearly 75% variant allele frequency (VAF) for the exon 19 deletion (ex19 DEL), indicating that deletion is present in most tumor cells, is plausible to hypothesize that the oncogenic transformation may have occurred in a common progenitor shared by AT2, AT1, or Clara cells. This possibility aligns with prior evidence suggesting that the founder mutation could arise in progenitor cells expressing members of the ERBB family(*26*) referred to as bronchioalveolar stem cells (BASCs), which have been implicated in the development of lung adenocarcinoma(*27*). Regarding the intriguingly dedifferentiated NE-like/EMT cell population, the absence of canonical neuroendocrine markers, namely INSM1, CHGA, and NCAM1 (CD56), together with NEUROD1, NCAM2, CDH4 expression and an overall EMT phenotype that lacks CD70, a marker associated with resistant EMT(*28*), suggests that this population represents a novel dedifferentiation trajectory induced by OSI treatment, still falling within the spectrum of SCLC subtypes(*29*). These results are timely considering recent findings on the EGFR × HER3 bispecific antibody, Iza-Bren, which targets both the EGFR and HER3 pathways and has shown encouraging efficacy, with a response rate of 75% in SCLC(*30*). Along these lines, this dual-targeted ADC therapy has demonstrated promising activity in a pooled analysis of phase 1 and phase 2 trials in EGFR-mutated NSCLC(*31*), and is currently being evaluated in a phase 3 trial in combination with OSI as first-line therapy (NCT06838273). Our data support the rationale of this trial and provide mechanistic insights into how the ERBB network, through EGFR/HER3 interactions, can reprogram tumor histology toward NE-like/EMT dedifferentiation. That a NE-like compartment produces NRG1 is probably not surprising, given that NRG1 is expressed almost exclusively in the tissue of neural origin where it plays a role in the neural regulation (*32*) of skeletal muscle differentiation and has been implicated as a neuron-induced differentiation agent in clonal neural crest cell cultures (*33, 34*). The immediate invasive potential of DTPs remains to be defined along with the pronounced capacity to infiltrate the extracellular matrix, consistent with an emergent metastatic phenotype. Whether this behaviour arises broadly across cancer cell clones or is restricted to those undergoing NE-like/EMT differentiation remains unresolved as well as a putative involvement of MYC and YAP pathway(*35*).

## Materials and Methods

### Experimental design

The objective of this study was to identify non-genomic mechanisms driving osimertinib tolerance in NSCLC drug-persistent cells. Besides PC9 tumor cell line we employed an arsenal of patient derived specimen treated with EGFR-TKIs. Patient samples were selected on the basis of tumor histology, drug treatment and the mutational status. We used quantitative real-time PCR, Western blotting, ELISA, droplet digital PCR, single-cell RNA sequencing and *in vivo* animal studies. Animals were randomly assigned to control or treatment groups. Investigators were not blinded to experiments. Additional details on cell lines, antibodies, and experimental procedures can be found in Supplementary Materials and Methods.

### Cell culture and patient samples

PC9 and ADK11LM were cultured in Roswell Park Memorial Institute (RPMI) 1640 medium containing 10% of fetal bovine serum (FBS) (Thermo Fisher Scientific), 1% penicillin/streptomycin (Corning Inc.) and 1% L-glutamine (Corning Inc.) in a 37°C atmosphere containing 5% CO2. ADK11LM early-DTPs were obtained from a malignant pleural effusion of an EGFR-mutated NSCLC patient with exon 19 deletion, who underwent progression with OSI. Cycling DTP (cDTP) cells (PC9 and ADL11LM) were maintained in culture with OSI at a concentration of 40nM. FFPE samples from patients (N=3) undergoing or progressing under TKI treatment were collected and RNA was extracted using RNeasy FFPE Kit (Qiagen #73504). The study received the approval by the Ethical Committee of the IRCCS Azienda Ospedaliero–Universitaria di Bologna (protocol no.591/2022/Sper/AOUBo).

### Colony-forming assay

800 cells/well of PC9 and 30.000 cells/well of ADK11LM were seeded in full media in 12-well plates and treated the following day according to the experiment. After 7 days, fresh treatments were added, and 10 days later, the cells were fixed in 4% paraformaldehyde (PFA) and stained with 0.5%. Crystal Violet. An image of each well was captured, and the covered area was measured using ImageJ software. For each treatment, the mean value of the covered area was calculated as a percentage of the control.

### 3D spheroid forming assay

For the spheroid formation assay 800 cells/well of PC9 and 30.000 cells/well of ADK11LM were grown in suspension over a layer of agar diluted to 0.6% in full medium. Treatments were added with the seeding. After 15 days, pictures of four non-overlapping fields for each well were collected. Spheroids in each picture were counted, and the size was quantified by measuring the length of the major and minor axis using ImageJ software. Axis values below 50 A.U. were excluded and volume was calculated through the following formula (major axis x minor axis^2^) /2.

### Immunofluorescence

For immunofluorescence, 50.000 cells were seeded in full medium onto glass coverslips and fixed the day after with 4% PFA. Then, cells were permeabilized with 0.2% Triton for 5’ and blocked for 1 hour in bovine serum albumin (BSA) 1%. An overnight incubation with primary antibodies (Ab I) was carried out in humidified chamber. A complete list of antibodies used is provided in Table 1. The day after Ab I incubation, glasses were incubated with fluorescent secondary antibodies for 1h, and then for 30’ with a mix of DAPI (Sigma Aldrich, St.Louise, Missouri, USA) and FITC-/TRITC conjugated Phalloidin (Sigma Aldrich, St.Louise, Missouri, USA). Imaging was performed with the Olympus BH-2 CCD microscope.

**Table 1.** Reagents. List of antibodies used for western blot, IF, IHC.

| ANTIBODY | SOURCE | CATALOG CODE |
| --- | --- | --- |
| **Western blot** |  |  |
| Phospho-AKT (Ser473) (D9E) XP Rabbit monoclonal antibody | Cell Signaling Technology | #4060 |
| AKT Rabbit polyclonal antibody | Cell Signaling Technology | #9272 |
| MAP Kinase activated (diphosphorylated ERK 1/2) Mouse monoclonal antibody | Sigma-Aldrich | #M8159 |
| ERK 2 (D-2) Mouse monoclonal antibody | Santa Cruz Biotechnology | #sc-1647 |
| EGFR (D38B1) rabbit monoclonal antibody | Cell signaling Technologies | #4267 |
| Phospho-EGFR (Tyr1068) (D7A5) rabbit monoclonal antibody | Cell signaling Technologies | #3777 |
| HER2 (D8F12) rabbit monoclonal antibody | Cell signaling Technologies | #4290 |
| Phospho-HER2 (Tyr1221/1222) (6B12) rabbit monoclonal antibody | Cell signaling Technologies | #2243 |
| HER3 (1B2E) rabbit monoclonal antibody | Cell Signaling Technology | #4754 |
| Phospho-HER3 (Tyr1289) (21D3) rabbit monoclonal antibody | Cell Signaling Technology | #4791 |
| GAPDH Rabbit antibody | Sigma-Aldrich | #G9545 |
| HER4 (111B2) rabbit monoclonal antibody | Cell signalling Technologies | #4795 |
| Phospho-HER4 (Tyr984) (Y984) rabbit monoclonal antibody | Cell signalling Technologies | #3790 |
| MET (D1C2) XP rabbit monoclonal antibody | Cell signalling Technologies | #8198 |
| Phospho-MET (Tyr1234/1235) (D26) XP rabbit monoclonal antibody | Cell Signaling Technology | #3077 |
| A-TUBULIN (B-7) mouse monoclonal antibody | Santa-Cruz Biotechnology | #sc-5286 |
| **Immunofluorescence** |  |  |
| Vimentin rabbit monoclonal antibody | Invitrogen | #MA5-14564 |
| EpCAM mouse monoclonal antibody | ABCAM | ab85987 |
| **Immunohistochemistry** |  |  |
| EpCAM mouse monoclonal antibody | Roche | #760-4383 |
| panCK-Anti-Keratin (34βE12) mouse monoclonal antibody | Roche | # 790-4373 |
| AE1/AE3 mouse monoclonal antibody | Roche | #760-2135 |
| anti-Thyroid Transcription Factor-1 (SP141) Rabbit Monoclonal Primary Antibody | Roche | # 790-4756 |
| anti-CD68 (KP-1)<br>Primary Antibody | Roche | # 790-2931 |
| CD138/syndecan-1 (B-A38) Mouse<br>Monoclonal Antibody | Roche | # 760-4248 |
| WT1 mouse<br>monoclonal antibody | Roche | #760-4397 |
| Vimentinanti-Vimentin<br>(V9) Primary Antibody | Roche | # 790-2917 |
| HER3 (1B2E) rabbit<br>monoclonal antibody<br>In vivo experiment | Cell Signaling Technology | #4754 |
| **Secondary antibodies**<br>**for**<br>**immunofluorescence** |  |  |
| Alexa Fluor® 488<br>AffiniPure Goat Anti-Rabbit IgG (H+L) | Jackson ImmunoResearch | #111-545-144 |
| Goat Anti-Mouse IgG<br>H&L<br>(Alexa Fluor® 647) | Abcam | #ab150115 |
| **Secondary antibodies**<br>**for Western blot** |  |  |
| Anti-Mouse/Anti-<br>Rabbit EnVision+<br>System- HRP Labelled<br>Polymer | Dako | #K4001/K4003 |

### Immunohistochemistry

Immunohistochemical staining for CKWS, AE1/AE3, EPCAM, TTF-1, CD68, NCAM and WT1 in either ADK11LM early-DTPs population or cycling DTPs ADK11LM was performed on a Benchmark Ultra automated system (Ventana Roche, Tucson, USA) using pre diluted antibodies. All specimens have been formalin-fixed and paraffin-embedded (FFPE) previously to staining. The antibodies used are reported in Table 1. The OptiView DAB IHC Detection Kit was used for detection. Immunohistochemical stainings for HER3 in xenograft tumors were performed as follows. All specimens have been formalin-fixed and paraffin-embedded (FFPE). Three-micrometer-thick sections were dewaxed and rehydrated. For antigen retrieval, sections were immersed in 200 ml citrate buffer (pH 6.0) or in tris-EDTA (pH 9.0) and heated in a microwave oven at 750 W for two 5-minute cycles. Slides were incubated with the primary antibody overnight at 4 °C (anti-HER3 (1B2E) rabbit monoclonal antibody (1:500, #4754 Cell Signaling Technology). The reaction was developed using a commercial streptavidine-biotine-peroxidase technique (ABC Kit Elite, Vector) and visualized with 3-amino-9-ethylcarbazole (Dako). Slides were counterstained with Harris hematoxylin.

### Transmission Electron Microscopy (TEM)

The ultrastructural features of eDTPs ADK11LM and cDTPs were examined with TEM. 30.000 cells/well were seeded in 6-well plate and cultured in incubator until confluence.

After washing, the cells were directly fixed in plate with 2.5% buffered glutaraldehyde at room temperature (RT) for 20 minutes and mechanically detached with scraper. Cells were recovered in microtubes, centrifuged and stored at 4°C overnight. After post-fixation in 1% buffered osmium tetroxide for 1 hours at RT, cellular pellets were dehydrated with ethanol (from 30% to 100%), passed in acetonitrile and resin embedded. Ultrathin sections (80 nm) were collected on grids and uranyl acetate staining followed by lead citrate before the observation in Philips CM10 (FEI Company, ThermoFisher, Waltham, MA, USA) Transmission Electron Microscope. Images were acquired thought an Olympus digital camera.

### Western blot

Cells were seeded in 6cm^2^ dishes, 1×10^6^ for PC9 and 2×10^6^ for ADK11LM. After seeding, cells were starved overnight and treated according to the experimental setting. For western blot, lysis was carried out with RIPA buffer supplemented with a protease inhibitor cocktail (P8340, Sigma-Aldrich, St. Louise, Missouri, USA, 1:100) and a phosphatase inhibitor solution containing of 1 mM Na3VO4 (Santa-Cruz Biotechnology, Dallas, USA). To measure protein concentration, DC Protein Assay (Bio-Rad, Hercules, USA) was employed and BSA was used as the standard. Protein lysates were then separated by SDS-PAGE electrophoresis and transferred to nitrocellulose membranes (Amersham™ Protran™ Premium, GE Healthcare, USA). Membranes were blocked for 1h with 3% BSA in TBS-T (0.1% Tween-20) before overnight incubation at 4°C with Ab I. Then, a 1h incubation with horseradish peroxidase-conjugated secondary antibodies (Ab II) was performed and detection was carried out by chemiluminescence (Amersham™ ECL™ Detection Reagents). A full list of antibodies used is provided in Table 1.

### RT-qPCR

1×10^6^ PC9 and 2×10^6^ ADK11LM cells were seeded in 6cm^2^ dishes in 3ml of 10% FBS medium and after 8 hours they were starved. The following day, treatments were added according to the experiments. Then, cells were washed with cold PBS 1X and incubated for 5 minutes at RT with 500 μL of QIAzol Lysis Reagent (Invitrogen, Life Technologies) to perform total RNA extraction from cells, according to manufacturer’s instructions. The extracted RNA was then converted into complementary DNA (cDNA). RNA reverse transcription (RT) was carried out using the High-Capacity cDNA Reverse Transcription Kit (Applied BiosystemsTM). RT-qPCR analyses were then performed on the cDNA using the SsoFast EvaGreen Supermix (Bio-Rad, Hercules, USA). Gene expression data from qRT-PCRs were determined as ΔCq (Quantification Cycle) normalized on the expression of β_2_-microglobulin (B2M) housekeeping gene. The primers used for qPCR are listed here Table 2.

**Table 2.**
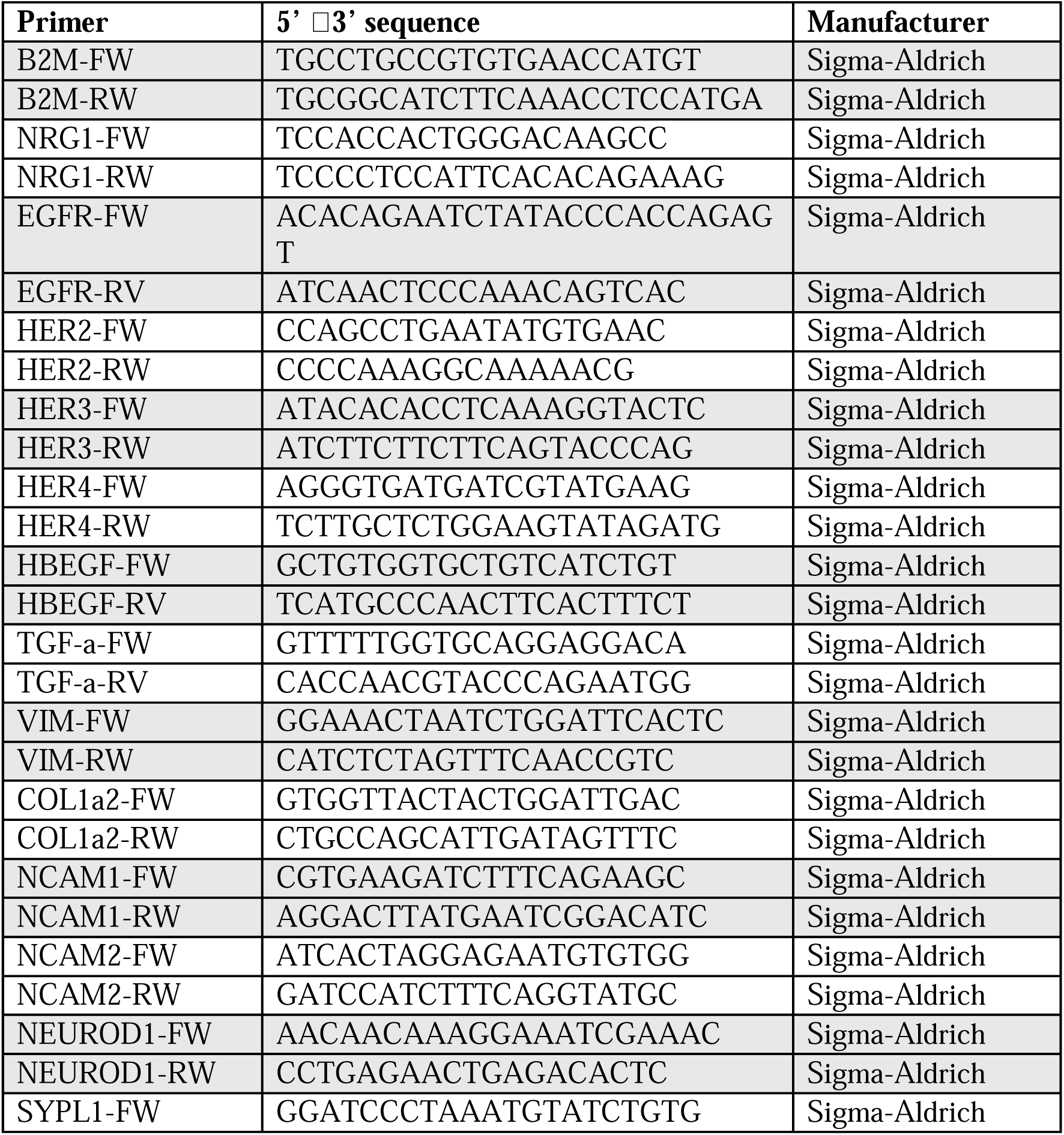
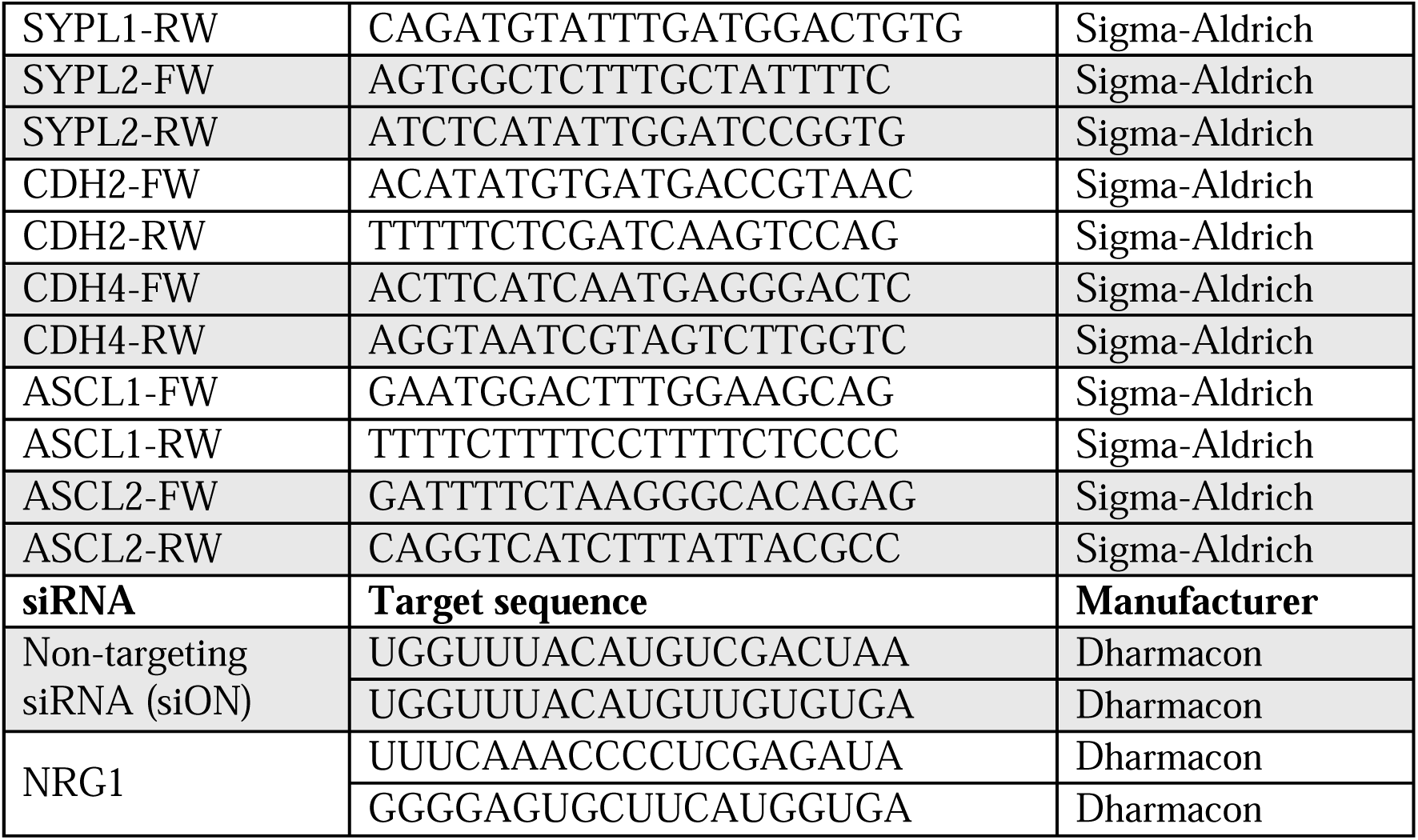
Reagents. List of primers used for RT-qPCR and siRNA.

### Droplet digital PCR (ddPCR)

Droplet Digital PCR (ddPCR) was performed using QX200 ddPCR EvaGreen SuperMix (Bio-Rad, Hercules, CA, USA, cat. no. 1864034) and QX200 Droplet Digital PCR System (Bio-Rad). Briefly, for each target gene a PCR mixture containing 10 μL of 2X QX200 ddPCR EvaGreen SuperMix, primers at final concentration of 250 nM, a variable volume input of cDNA in a final volume of 20 μL. All the experiments included a no template control (NTC). The cycling protocol was 95 °C for 5 min, then 40 cycles of 96 °C for 30 s and 58 °C for 1 min and three final steps at 4 °C for 5 min, 90 °C for 5 min and 4 °C infinite hold. The ramping rate between these steps was set at 2 °C/second. Data analysis was performed using the software QX Manager version 2.4 (Bio-Rad) and the positive droplets were manually selected using the specific tool. The final concentration of each target gene was obtained in all the analyzed samples, expressed in copies/μL. This concentration was normalized as copy number in 25 ng of cDNA.

### Single-cell and pathways enrichment analysis

Cells isolated from each population were captured in droplet emulsions using a Chromium Single-Cell instrument (10x Genomics) and libraries were prepared using the 10x Genomics 3′ Single Cell V2 protocol. All 10x libraries were pooled and sequenced on a NovaSeq 6000 (Illumina). For downstream analysis, data quality control, normalization, and dimensional reduction were performed using the R package Seurat (CRAN.R-project.org). We removed low-quality cells (e.g., cells with exceptionally low unique gene counts or abnormally high mitochondrial gene expression) and filtered out genes detected in very few cells only. Normalized data were then used for clustering to identify distinct cell populations, which were visualized using Uniform Manifold Approximation and Projection (UMAP). Cell types were assigned to each cluster using the computational platform ScType(*36*), a tool known for its robust cell type prediction. We augmented ScType’s default marker database with a set of manually curated, experiment-specific markers (see table S2) to refine our cell type annotations. To identify molecular changes, differential gene expression analysis was conducted between cell populations using Seurat’s built-in functions. Genes were considered differentially expressed if they exhibited a statistically significant change (adjusted p-value < 0.05) and a fold-change exceeding a defined threshold. The expression distribution of specific genes of interest was then visualized. Finally, to understand the biological implications of these transcriptional changes, pathway enrichment analysis was performed on the top 1000 overexpressed genes from the ‘Unknown’ clusters. This analysis utilized the ENRICHR database (amp.pharm.mssm.edu/Enrichr/), drawing upon established biological pathway and gene set resources (e.g., GO Biological Processes, KEGG pathways) to identify significantly enriched pathways (adjusted p-value < 0.05).

### Analysis of PC9 osimertinib time course

For the PC9 osimertinib time-course dataset, analyses were performed across days 0, 3, 7, and day 14 persisters, with the day 14 persister population stratified into cycling versus non-cycling subgroups as defined in the original study(*3*). Processed single-cell RNA-seq datasets were analyzed in Seurat (log-normalized expression). An NE-like/EMT transcriptional program score was computed per cell using Seurat’s AddModuleScore applied to a predefined EMT/NE marker gene set (table S3). Statistical analysis for group comparisons was performed versus day 0 using pairwise Wilcoxon rank-sum tests.

### Single-cell analysis of patient data

For the patient tumor and microenvironment analyses, cells were restricted to EGFR driver-gene patients and stratified by treatment stage: treatment-naive (naive), on-treatment drug-tolerant persisters (persisters), and relapse/progressive disease (relapse)(*13*). EMT_NE_like_score was calculated as previously described for PC9 cell line data. Pearson correlations between per-cell NRG1 expression and the EMT/NE-like module score (EMT_NE_like_score) were computed separately within each clinical group and cell type using per-cell values (NRG1 extracted with FetchData). Correlations were reported as NA when NRG1 showed no variation within a subgroup. Non-cancer epithelial cells were defined by inferCNV-based annotation (inferCNV_annotation = “nontumor”) and restricted to EGFR patients (driver_gene = “EGFR”). These non-tumor cells were subdivided into non-tumor fibroblasts versus non-tumor others using a fibroblast module score consisting of 9 genes (*COL1A1, COL1A2, COL5A1, LOXL1, LUM, FBLN1, FBLN2, CD34, PDGFRA*)(*37*) computed with Seurat AddModuleScore (). Cells were classified as non-tumor fibroblasts if their fibroblast module score exceeded the 95th percentile of the score distribution among validated non fibroblast cell types. All remaining non-tumor cells were classified as non-tumor others.

### NGS Mutation analysis

DNA was extracted with the QIAamp DNA Blood Mini QIAcube Kit (Qiagen, Hilden, Germany). QIAzol Lysis Reagent (Invitrogen, Life Technologies) was used to perform total RNA extraction from cells, according to manufacturer’s instructions. The DNA and RNA concentration was quantified using a Qubit fluorometer (Thermo Fisher Scientific, Waltham, MA, USA). The next-generation sequencing (NGS) analysis was performed using a fully automated NGS platform (Genexus, Thermo Fisher Scientifics) with the Oncomine Precision Assay (OPA) panel (Thermo Fisher Scientifics); this assay covers all the actionable genes in patients with NSCLC.

### siRNA transfection

For the siRNA transfection, 4×10^5^ PC9 and PC9 cDTP were seeded in 6-well plates with their growth medium without antibiotics. Twenty-four hours later, cells were transfected with siRNA by Lipofectamine 3000 (Invitrogen Co., Waltham, Massachusetts, USA) according to manufacturer’s instructions at a final concentration of 50nM. Twenty-four hours after transfection, cells were detached and re-seeded according to the experiment. Sequences of siRNA used are in Table 2. ON TARGETplus siRNAs were obtained from Dharmacon (Lafayette, CO, USA) and were used as control. siNRG1 was validated in (*38*).

### Gelatin degradation assay

Gelatin degradation assay was carried on by coating glass coverslips with Oregon green-488 conjugated gelatin from pig skin (Invitrogen^TM^, Waltham, USA) diluted to a concentration of 0.2 mg/mL in PBS + sucrose 2%. After coating, gelatin was treated for 15’ on ice with cold glutaraldehyde 0.5% in PBS and, subsequently, with NaBH_4_ 5mg/mL in PBS for 3’ at room temperature. Coated coverslips were stored at +4°C in PBS + P/S 1:50 protected from light until the seeding day. 50’000 cells were seeded and let attach for 6 hours. Cells were incubated for 24h with the different treatments. The coverslips were finally fixed with PFA 4% for 15’, blocked for 30’ with BSA3%+0.1 Triton X-100, and stained with Phalloidin-TRITC (Sigma-Aldrich, St. Louis, USA) and DAPI diluted in BSA 0.3%+0.1 Triton X-100 for 30’. Imaging was performed using the Olympus BH-2 CCD microscope. Images were analyzed using ImageJ software and degradation was measured in the green channel in terms of area of degraded gelatin (without green fluorescence) normalized on nuclei number (DAPI).

### Live-cell imaging – Phasefocus Livecyte^TM^

For Phasefocus Livecyte™ imaging, 2/3000 cells were seeded in full medium in 96-well plates and incubated overnight to allow cells’ adhesion before starting Livecyte™ monitoring. Images of cells were taken at 10X magnification every 2 hours for a total of 3 days. Proliferation and motility parameters were computed by Phasefocus Livecyte™ software.

### *In vivo* experiment

A total of 3×10^6^ PC9 per mouse has been injected subcutaneously in the right flank of female Foxn1nu athymic mice, aged six weeks. Mice harboring palpable tumors were divided into five different treatment groups: vehicle control, OSI (*5mg/kg*), OSI+CTX (5mg/Kg+0,2mg/mouse/injection), OSI+mAb anti-NRG1 (5mg/Kg+0,2mg/mouse/injection), OSI+CTX+mAb anti-NRG1(5mg/Kg + 0,1mg/mouse/injection for each antibody). Antibodies were injected intraperitoneally (IP) twice weekly, while Osimertinib was administered daily by oral gavage. Treatments continued until day 24 and tumor volume was monitored for the next month twice a week and calculated using the following formula *V* = 3.14 × (*W*^2^ × *L*)/6. Body weight was measured once a week. Mice were euthanized when tumor size reached 1,500 mm^3^. A parallel in vivo experiment was performed with three mice per treatment group, essentially as described above. Mice were sacrificed after seven days of treatment. The respective tumors were extracted and then processed for immunoblotting and immunohistochemistry using the indicated antibodies. Therapeutic Antibody antiNRG1 (YW538.24.71)(*39*) was acquired from Genentech through an MTA request (ID # OR-221436). Ethical approval for this experiment was granted by the Italian Ministry of Health (authorization no. 663/2022-PR).

### ELISA (enzyme linked immunosorit assay)

2×10^6^ cells were seeded in FM 10% FBS and starved after 8 hours. The day after the treatment was added accordingly with the experiment. Cell culturing supernatant was centrifuged at 500g, 4 °C, and collected to remove particulates. NRG1 was detected with a DuoSet® Human NRG1-β/HRG-β1 ELISA kit (DY377-05, R&D Systems) following the manufacturer’s instructions. The optical density (450nm subtracted by 560nm wavelength) was determined with a microplate reader (EnSpire, PerkinElmer) and a standard curve generated and used to calculate the NRG1 levels in the supernatant.

### Statistical analyses

The statistical analyses were performed by using Prism version 9 (GraphPad Software, Inc). Both T-test and one-way ANOVA were used to test significance of the assays. The details of the applied statistical test are reported in the figure legends. * Pvalue < 0.05; ** Pvalue < 0.01; *** Pvalue < 0.001; **** Pvalue < 0.0001.

## Funding

The research leading to these results has received funding by the AIRC IG 2024, Rif. 28760 to M.L. and from the Italian Ministry of Health, RC-2025-2794605 project to A.A. and M.L. Moreover, this research was supported by the Italian Ministry of University and Research under PRIN-PNRR2022 - to Mattia Lauriola. The views and opinions expressed are those of the authors only and do not necessarily reflect those of the European Union or the European Commission. Neither the European Union nor the European Commission can be held responsible for them.

## Author contributions

AM, CM, WK, IL, MS, MM, SV, GP, AP, performed and analyzed the *in vitro* experiments, execution, and interpretation of the assays. CG and DR carried out the *in vivo* studies in mice, including experimental procedures, data collection, and subsequent analysis. AM and ML were the major contributors in the design of the experiments and the writing of the manuscript, drafting and organizing the text. FA and MF carried out IHC staining and analysis. ADG, PLL, YO, YY, BG, AA and ML critically revised the manuscript, providing intellectual input and improving the clarity of the work. All authors read and approved the final version of the manuscript.

## Competing interests

The authors declare no competing interests

## Data and materials availability

All data generated or analyzed during this study are included in this published article [and its supplementary information files]

## Declaration of generative AI and AI-assisted technologies in the writing process

During the preparation of this work the authors used ChatGPT to improve language and readability. After using this tool, the authors reviewed and edited the content as needed and take full responsibility for the content of the publication.

## Notes

### Competing Interest Statement

The authors have declared no competing interest.

