## Supplementary Files for "Neuroendocrine-like/EMT dedifferentiation mediates resistance to EGFR inhibitors via the NRG1/HER3 axis"

Alessandra Morselli *et al.*

**This PDF file includes:**

Fig. S1 Immunochemistry evaluation of early-DTPs sample.

Fig. S2 Establishment of cycling-DTPs.

Fig. S3 scRNA-seq pathway enrichment NE-like/EMT persister cells and extended analyses of osimertinib-treated patients.

Fig. S4 Histopathological and molecular analyses in FFPE samples of osimertinib-treated patients.

Fig. S5 Extended analyses of the effects of NRG1 neutralization on invasion and proliferation of drug-tolerant persister cells.

Table S1 NGS analysis on sensitive and osimertinib persister NSCLC cell lines.

Table S2 List of genes used for cell clustering annotation.

Table S3 List of genes used for the NE/EMT score.

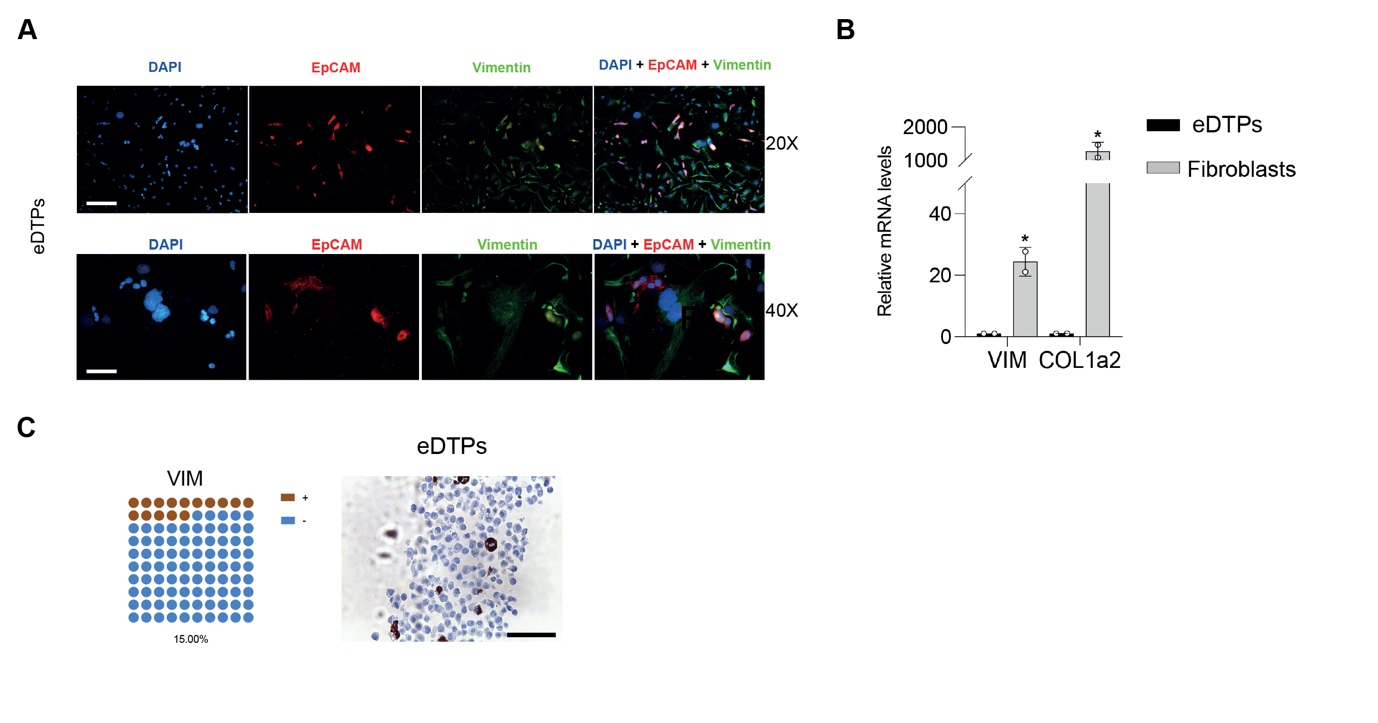
Fig. S1.

**Immunochemistry evaluation of early-DTPs sample.** (**A**) ADK11LM early -DTPs tumor sample stained with Vimentin (green), EpCAM (red) and counterstained with DAPI (blue). The magnification 20X (upper) and 40X (lower) are reported. Scale bar: 25µm and 50µm. (**B**) RNA was extracted from early-DTP tumor cells and fibroblasts derived from the same tumor sample. RT-qPCR was performed to analyze transcript levels of VIM and COL1a2. Data are reported as fold changes related to control ± SD. Unpaired T-test was applied for statistical analysis (**C**) IHC on ADK11LM early -DTPs tumor sample. The specimen was formalin-fixed and paraffin-embedded. The figure shows VIM IHC (20X magnification). Scale bar: 100 µm.

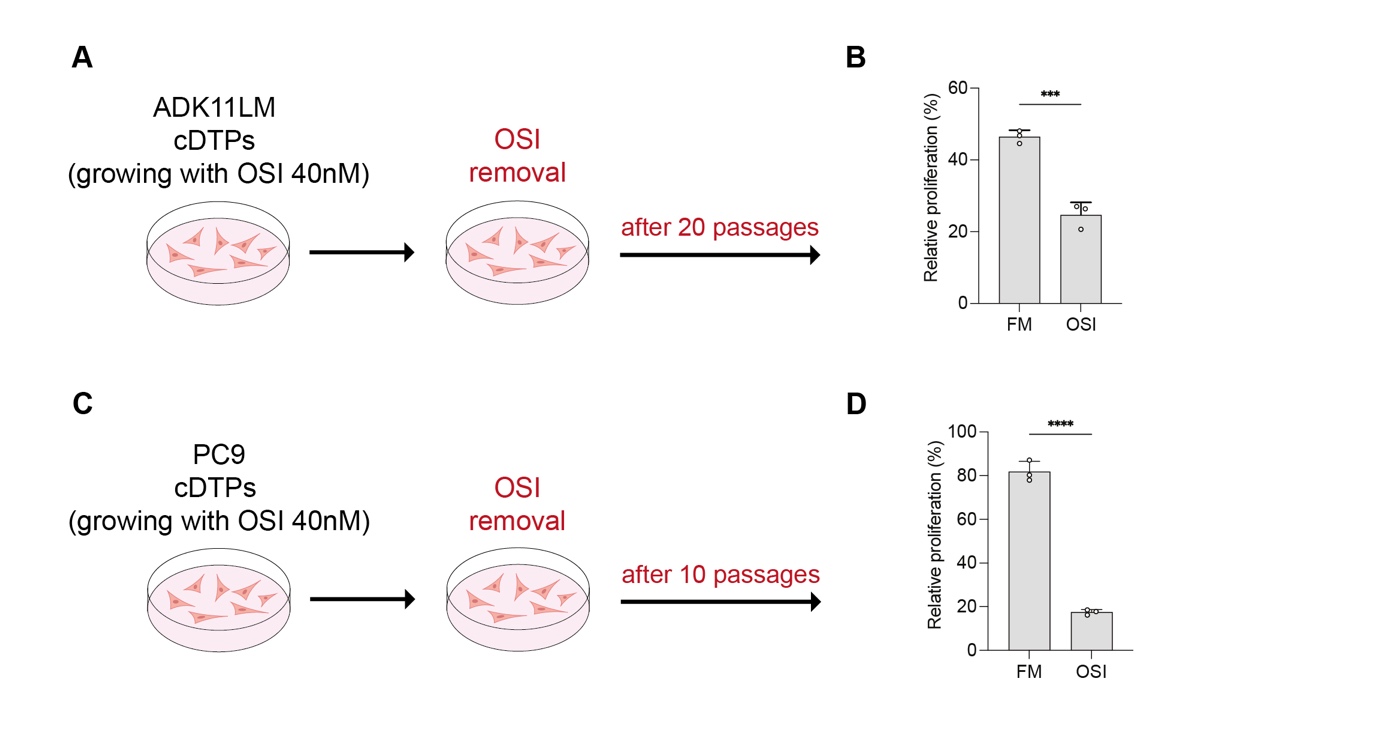
Fig. S2.

**Establishment of cycling-DTPs.** (**A**) ADK11LM cycling-DTPs were subjected to osimertinib washout and then allowed to grow in vitro. After 22 passages their sensitivity to osimertinib was tested. (**B**) Colony formation assay of ADK11LM cycling-DTP cells treated with, osimertinib (40nM) after 22 passages of drug holiday. Cells were seeded in triplicate in 12-well plates and treated after 24 hours. Following a 10-day incubation period, cells were fixed using 4% PFA and stained with crystal violet. Colony formation ability was quantified by counting the colonies using ImageJ software. Representative images of untreated and treated cells are shown. Data are reported as % of wells covered area in relation to controls ± SD. The statistic was calculated by one-way ANOVA test. (**C** and **D**) The same experiment was performed on PC9 cycling-DTPs.

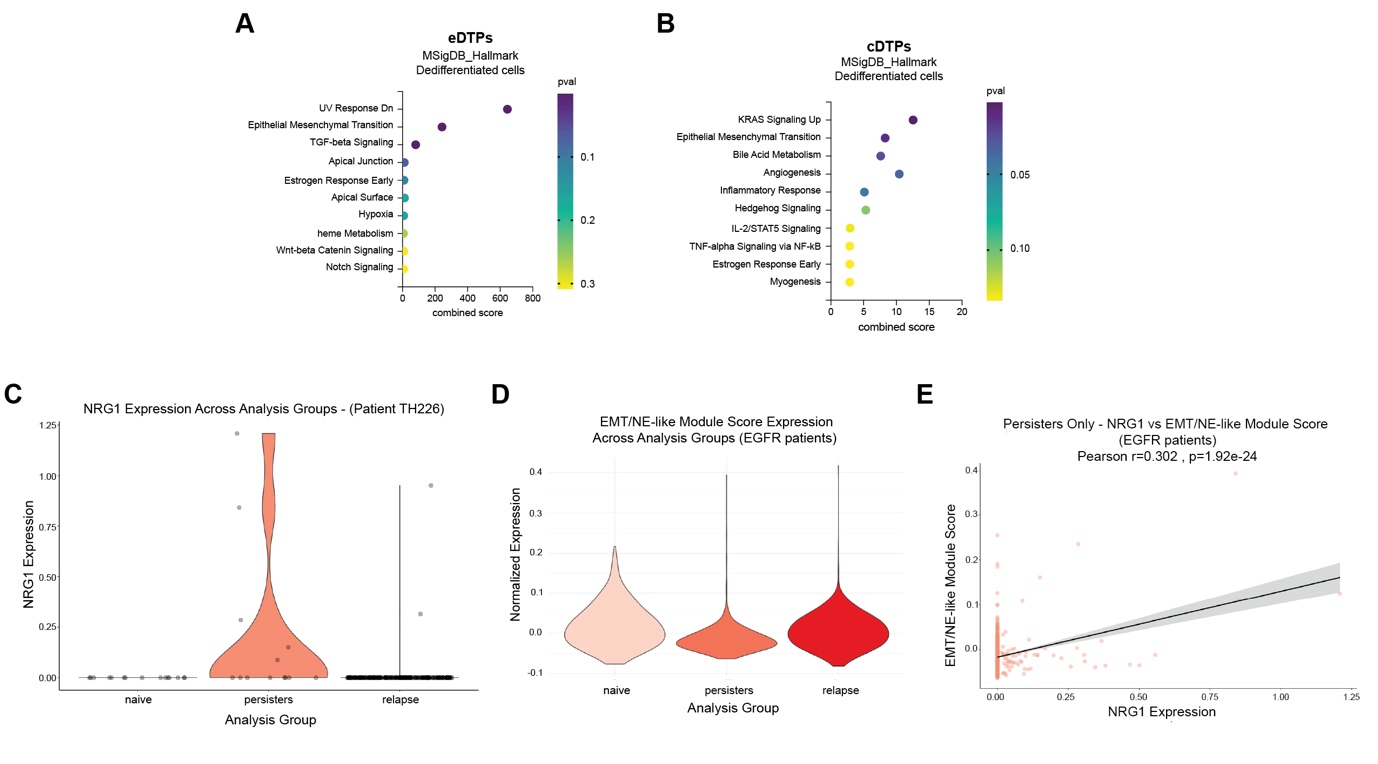
Fig. S3.

**scRNA-seq pathway enrichment NE-like/EMT persister cells and extended analyses of osimertinib-treated patients.** (**A** and **B**) Dot plot of pathway enrichment analysis performed on scRNAseq data showing the top 10 enriched MSigDB Hallmark with differential expression between the early and cycling -DTPs and the other cell types. The analysis was performed using the online tool Enrichr. Dot plot of pathway enrichment analysis performed on scRNAseq. Data showing the top 10 enriched Hallmark with differential expression between the NE-like/EMT early and cycling-DTPs and the other cell types. The analysis was performed using the online tool Enrichr. (**C**) NRG1 expression across treatment-naive (naive), persisters, and relapse cells in patient TH226 (Bivona dataset). Each dot represents a single cell. (**D**) NE-like/EMT module expression across EGFR-mutated NSCLC patients comparing naive, persisters, and relapse samples (Bivona dataset). (**E**) Relationship between NRG1 expression and EMT/NE-like module score in tumor persister cells.

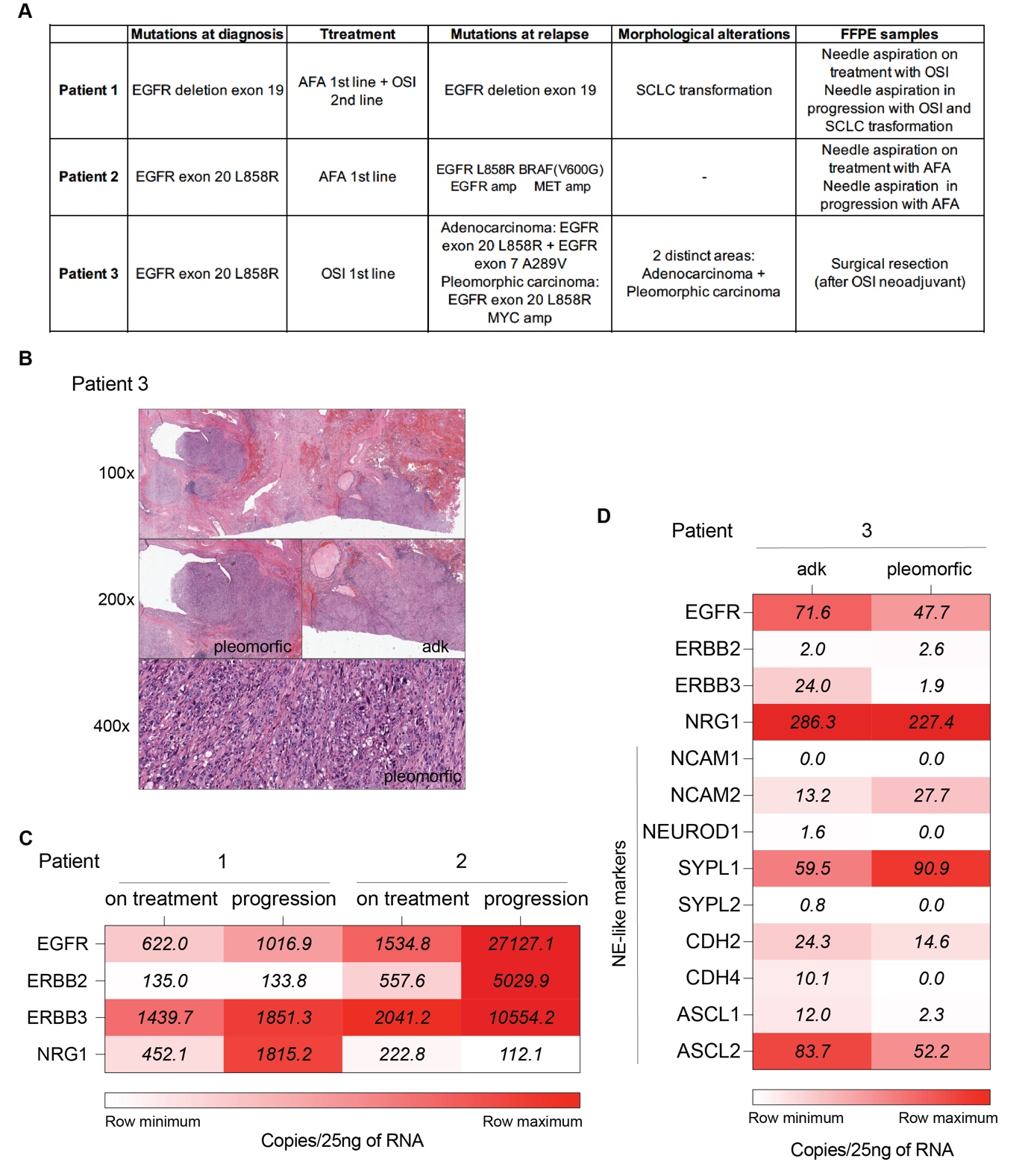
Fig. S4.

**Histopathological and molecular analyses in FFPE samples of osimertinib-treated patients.** (**A**) The table resume the clinical, histopatological and molecular features of patient 1, 2 and 3 derived specimens. (**B**) Histopathological features of a lung tumor resection specimen following neoadjuvant osimertinib.

The resection specimen shows a predominantly hilar tumor with marked intratumoral histomorphological heterogeneity. Top H&E, 100×: low-power examination reveals two spatially distinct components. On the right, a proliferation consistent with a nodal metastasis of lung adenocarcinoma with solid growth pattern, concordant with the pre-treatment biopsy diagnosis; on the left, a poorly differentiated carcinoma composed of spindle cells with marked nuclear pleomorphism, consistent with a pleomorphic carcinoma component.
left H&E, 200×: higher-power view of the solid adenocarcinoma component.
right H&E, 200× and bottom 400×: higher-power views of the pleomorphic component, highlighting spindle cell morphology and severe cytological atypia. Next-generation sequencing (NGS) identified an EGFR alteration in both components, while an additional MYC amplification was detected exclusively in the pleomorphic component, supporting molecular divergence potentially associated with treatment-related tumor evolution and aggressive histological transformation. (**C**) The heat map reports droplet-digital PCR performed on RNA extracted from FFPE samples belonging to patient 1 and 2. The expression of EGFR, ERBB2, ERBB3 and NRG1 is reported as copy number in 25 ng of RNA. (**D**) the same analysis was performed in patient 3 where RNA was extracted from the adenocarcinoma (adk) and pleomorphic component separately. The following transcript levels were analysed: EGFR, ERBB2, ERBB3, NRG1, NCAM1, NACM2, NEUROD1, SYPL1, SYPL2, CDG2, CDH4, ASCL1,ASCL2. their expression is reported as copy number in 25 ng of RNA.

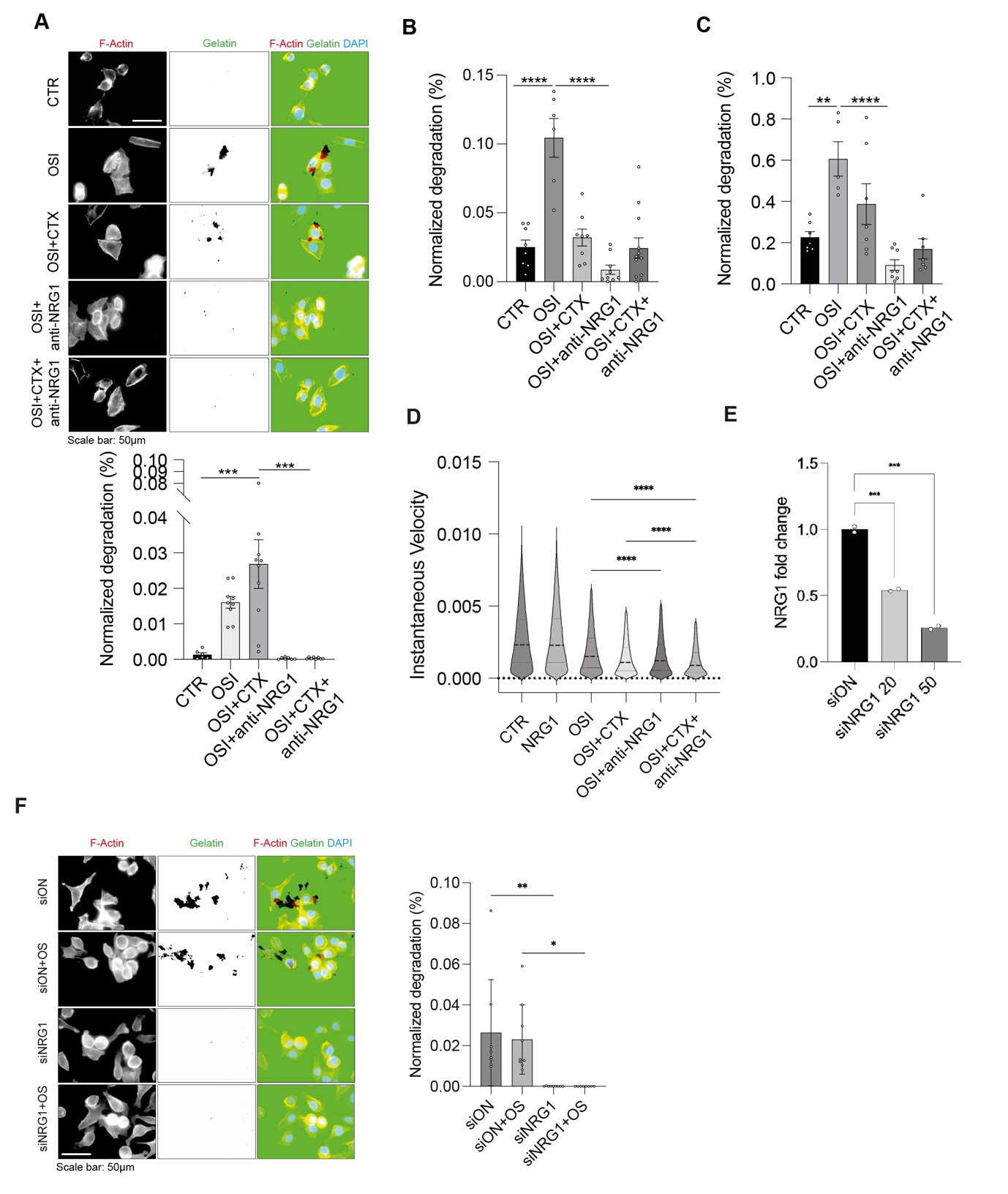
Fig. S5.

**Extended analyses of the effects of NRG1 neutralization on invasion and proliferation of drug-tolerant persister cells.** (**A**) Gelatin-degradation assay performed on PC9 cell line. The following treatments were applied: OSI (40nM), OSI + cetuximab (20 ug/mL), OSI + anti-NRG1 mAb (20 ug/mL) and the triple combination. When combined the doses of the two mAb were halved. After 24h, cells were stained with Phalloidin-TRITC (F-Actin) and DAPI (nuclei) prior to visualization. Invasion was quantified in terms of area of degraded gelatin (black spots visible in the green channel) normalized over the nuclei count in the same field. At least 7 fields were analyzed for each condition. Scale bar: 50 μm. (B and C) PC9 cycling-DTPs and ADK11LM cycling-DTPs were analyzed as in (**A**). (**D**) Quantification of PC9 instantaneous velocity over a 72h time-lapse obtained with the Phasefocus LiveCyte™ system, capturing images every 2h. Median and quartiles distribution are plotted. Outliers were identified and cleaned from results by means of ROUT method (Q = 1%). Statistic was calculated by one-way ANOVA test. Outliers were removed from the analysis. The statistic was calculated by One-way ANOVA. (**E**) Validation of NRG1 mRNA silencing through RT-qPCR. PC9 cells were transfected with the siRNA (20ng and 50ng) and the RNA was extracted. RT-qPCR was performed to analyze transcript levels of NRG1. Data are reported as fold changes related to control ± SD. One-way ANOVA test was applied. (**F**) Gelatin-degradation assay performed on PC9 cycling-DTP cells with a non-targeting control siRNA (siON) or siRNA targeting NRG1 (siNRG1). After siRNA transfection, cells were seeded and after 6 hours treated with osimertinib (40nM). After 24h, cells were stained with Phalloidin-TRITC (F-Actin) and DAPI (nuclei) prior to visualization. Invasion was quantified in terms of area of degraded gelatin (black spots visible in the green channel) normalized over the nuclei count in the same field. Scale bar: 50 μm. One-way ANOVA test was applied.

| Cell line | **Chromosome** | **Gene** | **Genic Region** | **Protein variant (cDNA)** | **Type of variant** | **Frequence** | **CNV(amplification)** |
| --- | --- | --- | --- | --- | --- | --- | --- |
| PC9 | 7 | EGFR | exon 19 | p.Glu746_Ala750del (c.2235 _2249del) | deletion | 88.4% | 7.2 |
| PC9 Cycling DTPs | 7 | EGFR | exon 19 | p.Glu746_Ala750del (c.2235 _2249del) | deletion | 11,40% | 49 |
| ADK11LM-early DTPs | 7 | EGFR | exon 19 | p.Leu747_Pro753delinsSer (c.2240_2257del) | delezione | 74,20% | - |
| ADK11LM Cycling DTPs | 7 | EGFR | exon 19 | p.Leu747_Pro753delinsSer (c.2240_2257del) | delezione | 74,20% | - |

**Table S1: NGS analysis on sensitive and osimertinib persister NSCLC cell lines.**

| **tissueType** | **cellName** | **geneSymbolmore** | **shortName** |
| --- | --- | --- | --- |
| Lung | Airway epithelial cells | ADH7,AQP1,CDH1,SEC14L3 | Airway epithelial |
| Lung | Airway goblet cells | AGR2,AQP5,CEACAM1,DMBT1,DUSP4,FXYD3,GALNT5,GALNT6,GGH,GOLPH3,GP2,IL19,LIPF,LTF,LYNX1,MSLN,MUC16,MUC4,MUC5B,NOS2,PIGR,QSOX1,SCGB3A1,SEC23B,SPDEF,TFF2 | Airway goblet |
| Lung | Mesothelial cells | C2,CALB2,CD44,CDH1,DES,EFEMP1,EPCAM,GSTA3,KRT5,KRT7,KRT72,LGALS7,LRRN4,ME1,ME2,MSLN,MUC16,OSR1,RSPO1,THBD,UPK3B,WT1 | Mesothelial |
| Lung | Fibroblasts | COL3A1,COL5A2,DPT,FN1,GSN,LRP1,PDGFRA,TCF21 | Fibroblasts |
| Lung | Basal cells (Airway progenitor cells) | ABI3BP,AQP3,DAPL1,GSTM2,HPGD,ICAM1,KRT14,KRT15,KRT5,PHLDA3,RPS18,SDC1 | Basal |
| Lung | Alveolar macrophages | ABCG1,CCL3,CD36,CLEC7A,CSF2RB,CXCL2,G0S2,GAL,GDA,GNGT2,GPNMB,IL18,IL1B,ITGAX,KLHDC4,MARCO,MCEMP1,MPP1,MRC1,OLR1,PLET1,S100A4,TLR2,TNFAIP2,TRIM25 | Alveolar macrophages |
| Lung | Ciliated cells | APPL2,ATP5MD,CCDC153,CCDC17,CCDC181,CCDC39,CYP2S1,FAM161A,FOXJ1,LRRC23,ODF3B,SCGB1A1,SEC14L3,SNTN,STK11,TMEM212,TSPAN19,TUBB4B,NEK10,SLC52A1,CDHR2,RAB36,POU2AF1,TSNAXIP1,CYP4X1,BICC1,KIF6,CLIC6,RP1,LPAR3,LRRC6,TMED6,ULK4,CAPSL,RFX2,ANKFN1,LPAR3,CCDC78,PROM1,HMGA2,WNT5B,TUB | Ciliated |
| Lung | EMT/Dedifferentiated cells | ZEB2,ZEB1,FBN1,COL21A1,ABCG2,ABCG5,ABCB1,TWIST1,UNC5,SOX9-AS1,SOX5,KLF15,TGFB2,IGF2BP1 | EMT |
| Lung | Club cells | LOXL4,CYP4X1,STRA6,LTF | Club |
| Lung | Clara cells | AHR,ALDH1A1,BPIFA1,CTSE,CYP2E1,CYP4B1,ERN1,FOXM1,LEPR,MUC1,MUC4,MUC5B,RAB3D,SCGB1A1,SCGB3A1,SCGB3A2,SFTPA1,SFTPC,SFTPD,SYT2,KYNU | Clara |
| Lung | Endothelial cell | CD34,EGFL7,EMCN,ESAM,FLT1,KDR,MCAM,PECAM1,RAMP2,TEK,VWF | Endothelial cell |
| Lung | Epithelial cells | ANPEP,EPCAM,IL10,IL6R | Epithelial |
| Lung | Ionocytes | CFTR,CLCNKB,FOXI1,KCNMA1,SCGB1A1,SLC12A2,TFCP2L1 | Ionocytes |
| Lung | Pulmonary alveolar type I cells | AGER,AKAP5,AQP3,AQP5,CCN2,CLDN18,CLIC5,COL4A3,COL4A4,CRLF1,CYP4B1,EGFL6,EMP2,FSTL3,GPRC5A,HOPX,ICAM1,IGFBP6,KRT7,MEX3B,MMP11,P2RX7,PDPN,PXDC1,RTKN2,SCNN1A,SCNN1B,SCNN1G,SEC14L3,SEMA3B,SEMA3E,SMARCA1,VEGFA,MST1R,IGFBPL1 | AT1 |
| Lung | Pulmonary alveolar type II cells | ABCA3,ADGRF5,AGER,CD36,CD3G,CEBPA,CLDN18,CRLF1,CTNND1,CXCL2,CXCR2,DDX3Y,EGFL6,ETV5,GRK2,IL1B,INMT,IRX1,LAMP3,LPCAT1,LRG1,MUC1,NAPSA,NKX2-1,NRN1,PGC,PIGR,PPBP,PPP1R14C,RUNX3,S100G,SDC1,SFTA2,SFTPA1,SFTPB,SFTPC,SFTPD,SLC34A2,SOAT1,CACNA2D2,LMNTD2 | AT2 |
| Lung | Secretory cell | MUC5B,PIGR,SCGB1A1,SLC4A4 | Secretory |
| Lung | Cancer cells | EPCAM,KRT9,KRT10,KRT12,KRT13,KRT14,KRT15,KRT16,KRT17,KRT18,KRT19,KRT20,KRT23,KRT24,KRT25,KRT26,KRT27,KRT28,KRT31,KRT32,KRT33A,KRT33B,KRT34,KRT35,KRT36,KRT37,KRT38,KRT39,KRT40,KRT1,KRT2,KRT3,KRT4,KRT5,KRT6A,KRT6B,KRT6C,KRT7,KRT8,KRT71,KRT72,KRT73,KRT74,KRT75,KRT76,KRT77,KRT78,KRT79,KRT80,KRT81,KRT82,KRT83,KRT84,KRT85,KRT86,SLC2A1,SLC7A11,EGFR,FGFBP2,LGR5,MYC,ERBB4,FGFR2,TGFB2,IGF2BP1 | Cancer |

**Table S2: List of genes used for cell clustering annotation.**

NE/EMT score

EMT / mesenchymal (22 genes)

TWIST1, TWIST2, ZEB1, SNAI3, MMP14, CD44, FBN1, COL3A1, COL4A2, COL5A1, COL5A2, COL6A2, COL6A3, COL7A1, COL12A1, COL16A1, PLOD2, POSTN, VCAN, CDH6, CDH11, FAP

NE / neuronal lineage (8 genes)

NCAM1, NCAM2, CDH2, CDH4, NEUROD1, SYPL2, PTH, OXTR

**Table S3: List of genes used for the NE/EMT score**
